# Long-range chemical signalling *in vivo* is regulated by mechanical signals

**DOI:** 10.1101/2024.02.15.580459

**Authors:** Eva K. Pillai, Sudipta Mukherjee, Niklas Gampl, Ross J. McGinn, Katrin A. Mooslehner, Julia M. Becker, Alex Winkel, Amelia J. Thompson, Kristian Franze

**Affiliations:** Department of Physiology, Development and Neuroscience, University of Cambridge, Downing Street, CB2 3DY, Cambridge, United Kingdom; Cell Biology and Biophysics Unit, European Molecular Biology Laboratory, Meyerhofstraße 1, 69117, Heidelberg, Germany; Developmental Biology Unit, European Molecular Biology Laboratory, Meyerhofstraße 1, 69117, Heidelberg, Germany; Institute of Medical Physics and Microtissue Engineering, Friedrich-Alexander-Universität Erlangen-Nürnberg, Kussmaulallee 2, 91054 Erlangen, Germany; Max-Planck-Zentrum für Physik und Medizin, Kussmaulallee 2, 91054 Erlangen, Germany; Wellcome-MRC Cambridge Stem Cell Institute, Jeffrey Cheah Biomedical Centre, Puddicombe Way, Cambridge Biomedical Campus, CB2 0AW

**Keywords:** axon pathfinding, neuronal guidance, mechanobiology, mechanotransduction, traction forces

## Abstract

Biological processes are regulated by chemical and mechanical signals, yet the interaction between these signalling modalities remains poorly understood. Using the developing *Xenopus laevis* brain as a model system, we identified a critical crosstalk between tissue stiffness and long-range chemical signalling *in vivo*. Targeted knockdown of the mechanosensitive ion channel Piezo1 in retinal ganglion cells (RGCs) led to pathfinding errors *in vivo.* However, pathfinding errors were also observed in RGCs expressing Piezo1, when Piezo1 was downregulated in the surrounding brain tissue. Depleting Piezo1 in the brain parenchyma led to a decrease in the expression of the long-range chemical guidance cues Semaphorin3A (Sema3A) and Slit1, which instruct turning responses in distant cells. Furthermore, Piezo1 knockdown markedly reduced tissue stiffness. This tissue softening was independent of Sema3A depletion, and was caused by a decrease in the cell-cell adhesion proteins NCAM1 and N-Cadherin. Downregulating NCAM1 and N-Cadherin was sufficient to reduce tissue stiffness and Sema3A expression. Conversely, increasing environmental stiffness *ex vivo* resulted in enhanced tissue-level force generation and an increase in Slit1 and Sema3A expression. Moreover, stiffening soft brain regions *in vivo* induced ectopic Sema3A production via a Piezo1-dependent mechanism. Hence, tissue mechanics can locally modulate the availability of diffusive, long-range chemical signals, thus influencing cell function at sites distant from the mechanical cue. Such indirect regulatory mechanisms of cell function through mechanical signals are likely widespread across biological systems.

## Introduction

Numerous biological processes are regulated by long-range concentration gradients of diffusible chemical signals, which act in concert to spatio-temporally control the function of cells within tissues^1^. Morphogen gradients, for example, set positional information during development^2–6^, while growth factor and hormone gradients are essential for organ development and maintenance^7^. Signalling molecules have multitudinous functions; for instance, the Semaphorin family of proteins regulate cell morphology and motility in the nervous, immune, respiratory, cardiovascular, and musculoskeletal systems during development, homeostasis, and disease^8^. Similarly, Slit proteins affect brain, kidney, heart, and mammary gland development, as well as in various disease states, by regulating cell proliferation, migration, vascularization and more^9^.

Recent work demonstrated that many of these cellular functions are also regulated by tissue mechanics. For instance, mechanical anisotropies in tissues affect growth orientation, while factors such as mechanical stresses, environmental stiffness, and cell or tissue geometry profoundly affect cell proliferation, migration, and fate specification^10–12^. In the nervous system, for example, mechanical forces regulate development and various physiological processes, while alterations in tissue mechanics occur in neurodegenerative diseases and other pathophysiological conditions^13^. Thus, besides the well-established role of chemical signals, mechanical cues are equally critical regulators of biological processes.

Indeed, cells will always be simultaneously exposed to both chemical and mechanical cues. One example is the extension of neuronal axons along precise paths during nervous system development, termed axon pathfinding. In the *Xenopus laevis* optic pathway, retinal ganglion cell (RGC) axons exhibit a stereotypical trajectory, exiting the eye via the optic nerve, crossing the midline at the optic chiasm, traversing the contralateral brain surface, and making a characteristic caudal turn at the mid-diencephalon before reaching their target, the optic tectum^14,15^ (Fig. 1a). Several membrane-bound and diffusible signalling molecules pattern the surrounding brain tissue, directing *Xenopus* RGC axons along this specific path^16^. For instance, the caudal turn of RGC axons in the mid-diencephalon is regulated by diffusive gradients of the repulsive long-range guidance cues, Slit1^17^ and Semaphorin3A (Sema3A)^18^. At the same time, RGC axons are mechanosensitive and encounter an instructive stiffness gradient in the brain, which also contributes to regulating RGC axon pathfinding in that region^19,20^. However, little is currently known about how cells integrate chemical and mechanical signalling *in vivo*. While recent studies exquisitely showed how environmental mechanics may regulate cell-intrinsic gene expression patterns and chemical signalling pathways^21–24^, how mechanical cues influence cell-extrinsic signalling—and, consequently, the function of distant cells not directly exposed to the mechanical signal—remains elusive.

**Figure 1:**
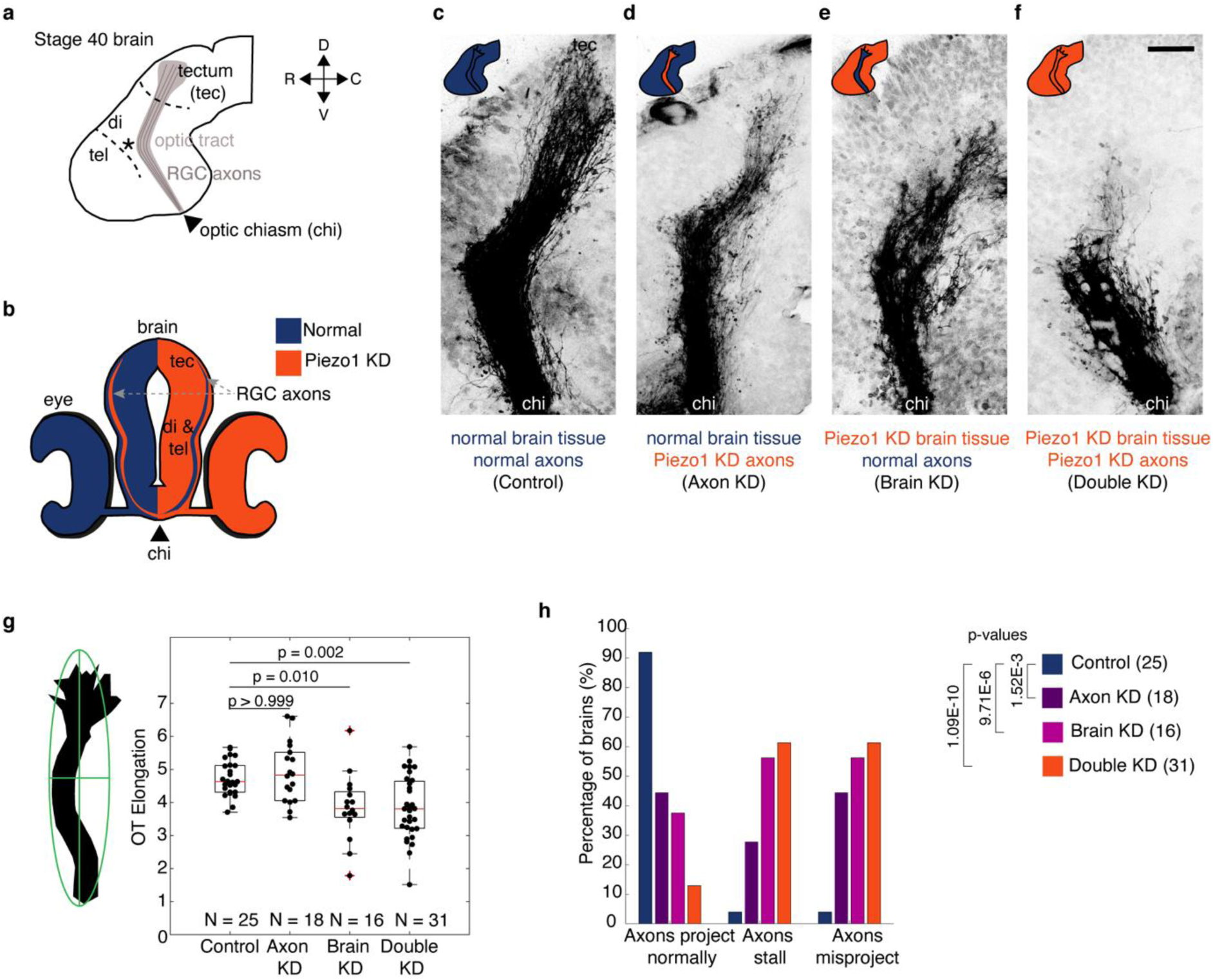
RGC intrinsic and extrinsic Piezo1 signalling is required for accurate axon pathfinding *in vivo*. **a,** Schematic of the lateral view of a stage 40 *Xenopus* brain with orientation guides for Rostral-Caudal (R, C) and Dorsal-Ventral (D,V) axes. RGC axons grow from the optic chiasm towards the optic tectum along a stereotypic path (the optic tract), turning caudally at the mid-diencephalon (“*”). **b,** Schematic cross-section of a Xenopus brain and retinae when Piezo1 is downregulated unilaterally in the nervous system. RGC axons cross the midline at the optic chiasm and grow across the contralateral brain surface. Normal Piezo1 levels are indicated in blue while Piezo1 depletion is shown in red. Piezo1- depleted axons grow across brain tissue with normal Piezo1 levels and *vice-versa*. **c-f,** Images of RGC axon growth *in vivo*. Axon growth and guidance were affected by downregulating Piezo1 (**d)** in the axons, (**e**) in the surrounding brain tissue, or (**f**) in both. Scale bar: 50 μm. **g,** Optic tract elongation, the ratio of long to short axes on a fitted ellipse (schematic), decreased significantly when Piezo1 was downregulated in the brain parenchyma. Each point represents a brain. Boxplots show median, first, and third quartiles; whiskers show the spread of data; ‘+’ indicate outliers. N denotes the number of animals. Data were assessed with a Kruskal-Wallis test, p < 0.0001, and with a Dunn post-hoc test (adjusted p-values provided in the figure). **h,** Scoring of brains displaying aberrant phenotypes. Knocking down Piezo1 solely in the axons or brain tissue, or in both, result in significant stalling/misprojection defects (*p =* 1.014E-8; Chi-Squared test followed by Fisher’s Exact pot-hoc tests, number of animals in parentheses), data were from a minimum of three independent experiments. chi: chiasm, di: diencephalon, KD: knockdown,, RGC: retinal ganglion cell, tec: tectum, tel: telencephalon.

## Results

### Retinal ganglion cell-intrinsic and extrinsic signalling via Piezo1 is required for accurate axon pathfinding in vivo

A key player involved in sensing and responding to brain tissue stiffness^19,25^ is the mechanosensitive nonselective cation channel Piezo1^26,27^. Piezo1 is found in a broad range of animal tissues including the bladder, kidneys, lung, and skin^26^, as well as in the developing *Xenopus* nervous system^19,28^. Downregulating Piezo1 expression in the nervous system *in vivo* leads to aberrant axon growth and severe pathfinding defects during development^19^.

To disentangle the effect of Piezo1 activity in RGC axons from that of the surrounding neuroepithelial cells on axon pathfinding, we selectively downregulated Piezo1 in the axons while preserving its expression in the surrounding brain tissue.

Classical fate mapping experiments have shown that the 2 dorsal blastomeres of 4 cell-stage embryos contribute to the formation of the nervous system, with each blastomere contributing to one half of it^29,30^. Injecting translation-blocking morpholinos into one of the two dorsal blastomeres at the 4-cell stage (Supplementary Fig. 1a) therefore resulted in embryos with Piezo1-depletion in one half of the nervous system. In pre-metamorphic *Xenopus* embryos, all RGC axons cross the midline at the optic chiasm to the contralateral surface, without forming ipsilateral projections^31^ (which are typically only formed from Stage 52^32^). Thus, this approach yielded embryos with Piezo1-depleted RGC axons growing across healthy brain tissue, while in the contralateral brain hemisphere, normal RGC axons grew across Piezo1-depleted brain tissue (Fig. 1b, Supplementary Fig. 1). While it was challenging to confirm Piezo1 depletion specifically in RGC axons *in vivo*, as Piezo1 is expressed not only in the axons^19^ but also ubiquitously throughout the surrounding brain parenchyma, local Piezo1 depletion was clearly visible in the parenchyma of the tissue arising from the injected blastomere (Supplementary Fig. 1).

RGC axons were labelled with DiI, and their trajectories analysed at developmental stage 40^33^, when axons had reached their end target, the optic tectum. Consistent with our previous findings^19^, simultaneous downregulation of Piezo1 in both axons and the surrounding brain tissue led to severe axon pathfinding defects (Fig. 1c,f), compared with age-matched sibling embryos, where both dorsal blastomeres were injected with a control morpholino. The elongation of the RGC axon bundle growing along the brain surface, or optic tract, was significantly lower than in controls (*p =* 0.002, Kruskal-Wallis with Dunn post-hoc test) (Fig. 1g). Furthermore, only 13% of brains had normal axonal projections, while 61% of brains exhibited stalling and/or misprojection defects (i.e., deviations from the normal trajectory, such as bypassing the caudal turn in the mid-diencephalon or misrouting dorso-anteriorly into the telencephalon) (Fig. 1h). Representative images illustrating normal RGC axonal projections and various axon guidance defects are shown in Supplementary Fig. 2.

When Piezo1 was selectively depleted in RGC axons (Fig. 1d), optic tract elongation was similar as in controls (p > 0.999, Kruskal-Wallis with Dunn post-hoc test, Fig. 1g).

Nonetheless, only 44% of these brains exhibited normal axonal projections (Fig. 1h). Piezo1 downregulation in the axons resulted in 28% of brains with stalling defects, while 44% of the brains showed misprojecting or both stalling and misprojecting defects (Fig. 1h). These results confirmed that RGC axons require Piezo1 to respond to their mechanical environment, and that this cell-intrinsic mechanosensing is necessary for accurate pathfinding. However, the overall phenotype was milder than what we had observed in brains with downregulated Piezo1 expression in both RGC axons and the surrounding neuroepithelium^19^ (Fig. 1f-h), suggesting that Piezo1 expression in neuroepithelial cells was also required for proper axon pathfinding.

Indeed, when Piezo1 downregulation was limited to the surrounding brain tissue, axon pathfinding of healthy RGCs containing Piezo1 was also perturbed. In fact, the observed guidance defects were even more profound than in brains with only axonal Piezo1 knockdown (Fig. 1e). Optic tract elongation was significantly decreased when compared to controls (*p =* 0.010, Kruskal-Wallis with Dunn post-hoc test) (Fig. 1g), and only 38% of brains showed normal axonal projections. In contrast, 56% of brains exhibited stalling and/or splaying (i.e., unbundling) and misprojecting axons (Fig. 1h). Thus, despite normal Piezo1 levels in RGC axons, axon growth patterns were disrupted, confirming that Piezo1-mediated mechanosensing by neuroepithelial cells was also essential for accurate axon pathfinding.

### Piezo1-dependent expression of long-range chemical guidance cues in vivo

We next investigated how perturbations of neuroepithelial mechanosensing may lead to disruptions in growth patterns of healthy axons. At least two RGC-extrinsic factors are crucial for the turning of the optic tract in the mid-diencephalon (Fig. 1a): local tissue stiffness^19^ and long-range chemical signalling by the repulsive guidance cues Slit1^17^ and Semaphorin3A (Sema3A)^18^, which are both located rostrally to the optic tract’s caudal turn (Fig. 2a,e). Chemical perturbations of brain tissue stiffness *in vivo* mostly lead to a splaying of RGC axons in the diencephalon^19^, while depletion of Slit1 and Sema3A signalling predominantly leads to a stalled axon phenotype^17,34^. Since we found both axon guidance defects after downregulating Piezo1 expression in neuroepithelial cells (Fig. 1e,h), we first tested the effect of Piezo1 downregulation on the expression of these chemical guidance cues using hybridization chain reaction-fluorescence in situ hybridization (HCR-FISH).

**Figure 2:**
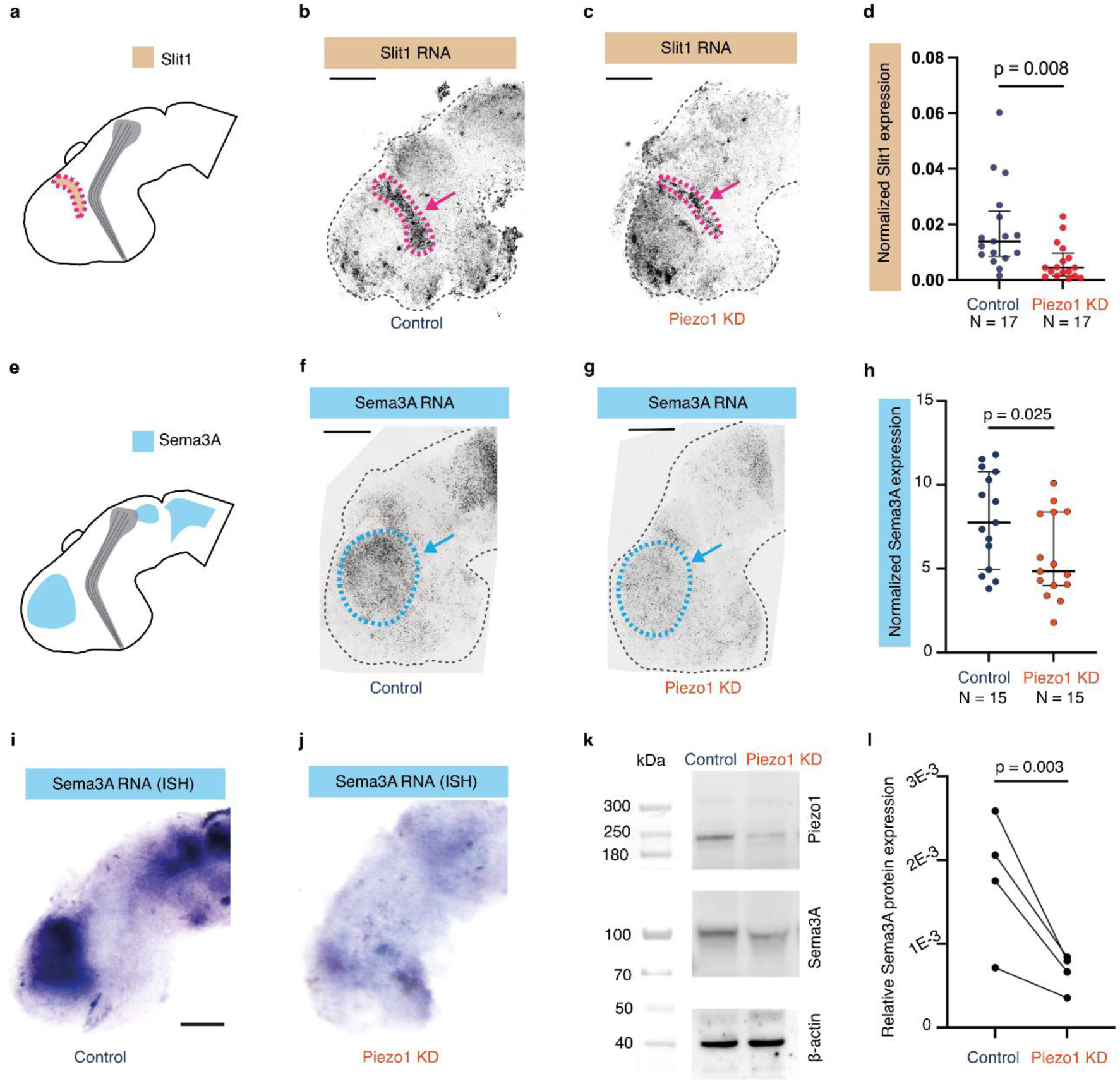
Piezo1 downregulation attenuates the expression of long-range chemical guidance cues *in vivo*. **a,** Schematic representation of the expression pattern of the chemical guidance cue, Slit1. **b-c,** Representative hybridization chain reaction-fluorescence *in situ* hybridisation (HCR-FISH) images of Slit1 expression in (**b**) a control and (**c**) a Piezo1 knockdown brain. **d,** Quantification of Slit1 mRNA expression. Normalised Slit1 expression decreased significantly in Piezo1 knockdown brains compared to control brains. **e,** Schematic of the expression pattern of Sema3A. **f-g,** Representative HCR-FISH images of Sema3A expression in (**f**) a control and (**g**) in a Piezo1 knockdown brain. **h,** Quantification of normalized Sema3A expression. Sema3A expression decreased significantly when Piezo1 was downregulated. **i-j,** *In situ* hybridisation of Sema3A mRNA expression, confirming a dramatic attenuation of the Sema3A signal in Piezo1 knockdown brains (**j**) compared to controls (**i**). **k,** Western blot image of Piezo1, Sema3A and β-actin protein expression in control and Piezo1-depleted brains. **l,** Western blot quantification. Relative protein expression levels were normalised to total protein staining. Downregulating Piezo1 led to significantly decreased Sema3A protein expression. Each point in **d** & **h** represents an embryo, lower quartile, median, and upper quartiles are indicated. N denotes the number of animals. Data in **d** & **h** were assessed with an unpaired t-test with Welch’s correction. Each point in **l** indicates the mean value of a biological replicate. Data in **l** was assessed with a ratio paired t-test. p-values shown in the figures. Scale bars: 100 μm. ISH: *In situ* hybridisation, KD: knockdown.

In stage 40 control-injected sibling embryos, our HCR-FISH results recapitulate published *in situ* hybridization data^17,18,35^: Slit1 forms a distinct band between the telencephalon and diencephalon, while Sema3A is highly expressed rostro-ventrally of Slit1 in the telencephalon (Fig. 2b,f). Piezo1 downregulation resulted in a significant reduction in Slit1 mRNA expression (*p =* 0.008, unpaired t-test with Welch’s correction) (Fig. 2c,d), and Sema3A mRNA expression was significantly decreased compared to controls (*p =* 0.025, unpaired t-test with Welch’s correction) (Fig. 2g,h). Standard *in situ* hybridizations confirmed the dramatic decrease in Sema3A mRNA expression in Piezo1-depleted brain tissue (Fig. 2 i,j). Additionally, Western blot analysis confirmed that Piezo1 downregulation resulted in significant decreases in Sema3A (*p =* 0.003, ratio paired t-test) (Fig. 2k,l) and Piezo1 protein levels, but not in the housekeeping protein β-actin, (Fig. 2k and Supplementary Fig. 3); Slit1 protein expression could not be assessed due to a lack of suitable antibodies in *Xenopus*. Hence, our data indicated that the downregulation of Piezo1 – a mechanosensor – in neuroepithelial cells attenuated the expression of long-range chemical cues, implying a role for mechanical signalling in modulating chemical signalling during development.

### Piezo1 depletion, but not Sema3A downregulation, leads to brain tissue softening

The splaying of axons seen in Piezo1-downregulated brain parenchyma (Fig. 1e,f) recapitulated the phenotype of RGC axons growing through brain tissue that was chemically softened *in vivo*^19^. To test if, in addition to the chemical landscape, tissue mechanics was also altered in Piezo1 knockdown brains, we employed atomic force microscopy (AFM)- based stiffness mapping^19,20^ to measure the stiffness of live developing *Xenopus* brains at stage 40. Tissue stiffness was quantified by the reduced apparent elastic modulus, *K*, whereby a larger *K* value indicates stiffer tissue. Measurements were performed *in vivo* in a rectangular grid on the exposed brain surface (Fig. 3a,b) of stage-matched sibling embryos for control injected embryos and embryos in which Piezo1 was downregulated either exclusively in the axons or in the surrounding brain tissue, or in both. We have previously shown that RGC axons do not contribute significantly to brain tissue stiffness^19^; thus, variations in axon growth are unlikely to affect the measured stiffness here.

**Figure 3:**
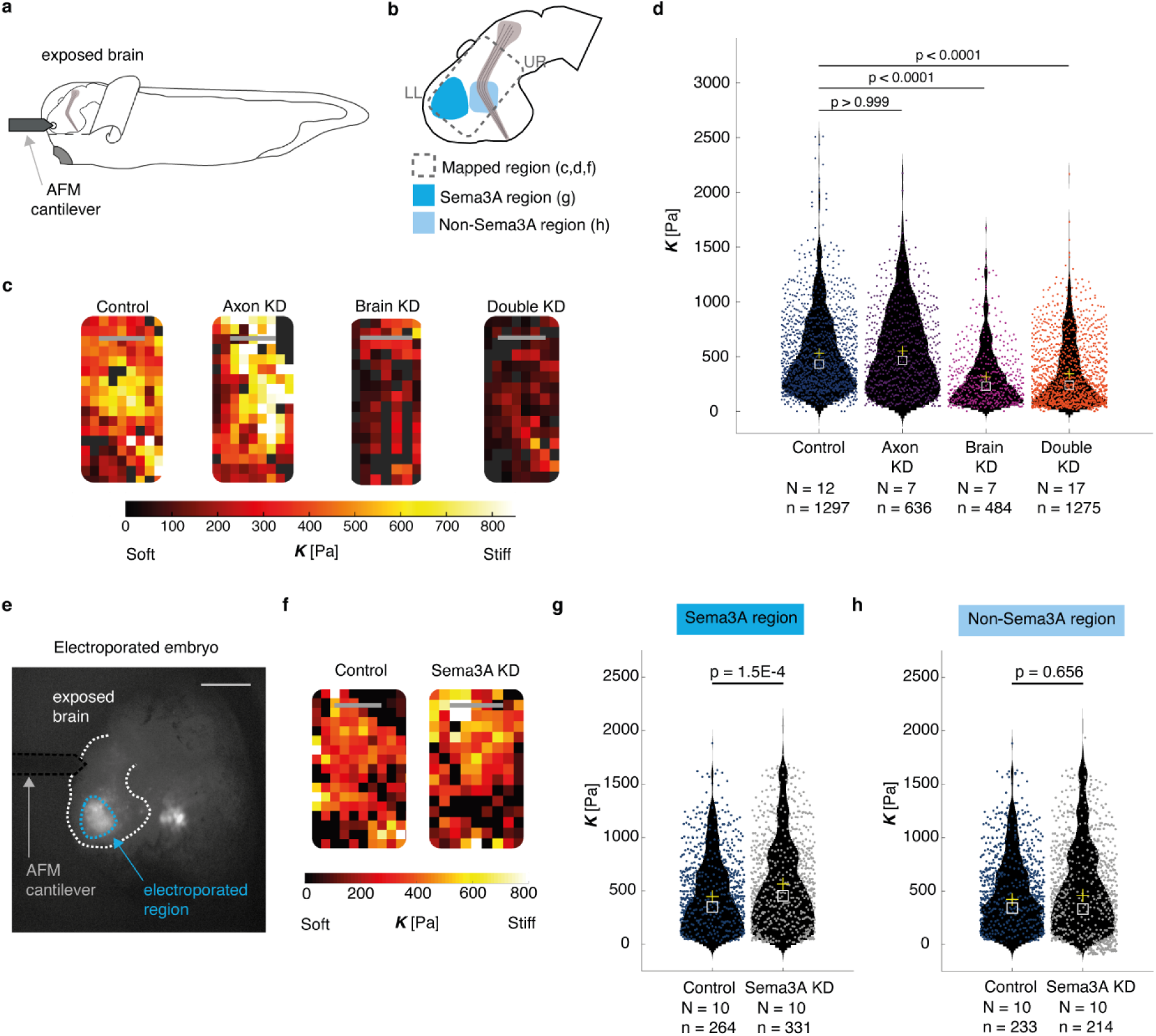
Piezo1, but not Sema3A, knockdown leads to brain tissue softening. **a,** Schematic of the experimental setup for *in vivo* brain-stiffness mapping. **b,** *Xenopus* brain schematics. The dashed rectangle indicates the mapped region, lower left (LL) and upper right (UR) corners of stiffness maps (relevant to **c** and **f**) are indicated, the colours indicate areas selected for regional analysis (relevant to **g** and **h**). **c,** AFM-based stiffness maps (colour maps), encoding the apparent elastic modulus, *K*, a measure of tissue stiffness, assessed at an indentation force *F* = 10 nN for control or Piezo1 downregulation exclusively in axons, in the surrounding brain tissue, or both. **d,** Quantification of AFM experiments. Downregulating Piezo1 in the brain parenchyma resulted in significant tissue softening. Violin plots with scatter of individual stiffness measurements for each condition. The means and medians are indicated as yellow crosses and white squares, respectively. Data was assessed with a Kruskal-Wallis test, p < 0.0001, followed by Dunn post-hoc tests (adjusted p-values provided in the figure). **e,** Exposed brain preparation of a stage 40 *Xenopus* embryo electroporated with fluorescein-tagged morpholinos, enabling the visualisation of the electroporated region. The dashed lines show the outline of the brain (white), the electroporated region (blue), and the AFM cantilever (black). Scale bar: 250 µm. (**f-h**) Downregulating Sema3A does not cause brain tissue softening. **f,** AFM-based stiffness maps (colour maps) for control or Sema3A morpholino-electroporated brains. **g-h,** Violin plots with scatter of individual stiffness measurements for each condition. Data were assessed with a Wilcoxon rank sum test. **g,** The electroporated regions were significantly stiffer in the Sema3A KD compared with the controls whereas **h,** the adjacent regions were mechanically similar. Scale bars: 100 μm. N denotes the number of animals, n denotes the number of measurements. KD: knockdown.

The overall mechanical landscape of brains with Piezo1-downregulation in solely the RGC axons was similar to that of control embryos (median *Kctrl* = 432 Pa, *KAxon k/d* = 467 Pa; p > 0.999, Kruskal-Wallis with Dunn post-hoc test) (Fig. 3c,d), suggesting that the observed axon pathfinding defects shown in Fig. 1d were not a consequence of alterations in tissue stiffness but rather originated from the axons’ reduced ability to sense their mechanical environment. Conversely, downregulating Piezo1 either exclusively in the surrounding brain tissue or simultaneously in the axons and surrounding brain tissue resulted in a close to two-fold decrease in brain stiffness (median *KBrain k/d* = 231 Pa, *KDouble k/d* = 240 Pa; p < 0.0001, Kruskal-Wallis with Dunn post-hoc test) (Fig. 3c,d). Hence, the knockdown of Piezo1 in the developing neuroepithelium led to considerable changes in both the chemical and mechanical landscape encountered by growing RGC axons.

In order to test whether the observed tissue softening following knockdown of Piezo1 was specific to the central nervous system or a more general phenomenon, we also measured the stiffness of skin in *Xenopus* embryos at stages 28-31 using AFM. Here too, Piezo1 knockdown led to a significant softening of the tissue (Supplementary Fig. 4), indicating that Piezo1 may not only sense but also regulate tissue stiffness in other organ systems.

We then wanted to know whether there was a causal relationship between the decrease in guidance cue expression (Fig. 2) and the decrease in tissue stiffness (Fig. 3c,d) in Piezo1 knockdown brains. In particular, we asked if tissue softening was a consequence of the decrease in Sema3A and Slit1, or if, vice versa, the decrease in guidance cue expression was a consequence of the change in tissue stiffness. Given the spatially restricted expression of Slit1 mRNA and the uncertainty if protein levels had changed after Piezo1 knockdown, we here focused on the effect of Sema3A depletion on tissue stiffness.

To downregulate endogenous Sema3A protein in the brain without altering the early developmental expression of Piezo1 levels, we electroporated the forebrain of stage 29/30 embryos with either a Sema3A or a control fluorescein-tagged morpholino^34^. The electroporated embryos were allowed to develop to stage 40, at which point tissue stiffness was measured using AFM (Fig. 3e). Downregulating Sema3A protein expression in the forebrain did not result in brain tissue softening. In fact, the stiffness of the Sema3A-depleted forebrain region increased compared to controls, with a median *K* = 455 Pa in the Sema3A knockdown brain and *K* = 350 Pa in the control (p < 0.0001, Wilcoxon rank-sum test) (Fig. 3f,g), while brain tissue mechanics in adjacent regions was unaltered in these embryos (median *KSema3A k/d* = 332 Pa, *Kctrl* = 341 Pa; *p =* 0.67, Wilcoxon rank-sum test) (Fig 3h). These data showed that the observed stiffening was specific to the localized Sema3A depletion, and suggested that the decrease in guidance cues in Piezo1 knockdown brains is likely not responsible for the decrease in tissue stiffness.

### Piezo1-mediated brain tissue softening occurs via depletion of cell-cell adhesion

The mechanical properties of brain tissue depend on its building blocks and their connectivity^36^. To understand why tissue softens following Piezo1 downregulation, we first assessed whether Piezo1 knockdown had an effect on cell body densities in the tissue, which are a well-established contributor to tissue stiffness in the developing *Xenopus* brain^19,20^. In agreement with previous literature^19,20^, in control embryos the density of cells rostral to the optic tract was higher than the density caudal to it (Fig. 4a-b). Rostral and caudal cell densities were unaffected upon either exclusively downregulating Piezo1 in the optic tract or the developing brain, or downregulating Piezo1 in both the brain and the optic tract (*p*rostral = 0.367 and *p*caudal = 0.454, One-way ANOVA) (Fig. 4b), indicating that the softening of the tissue following Piezo1 depletion could not be explained by changes in cell body densities.

**Fig. 4:**
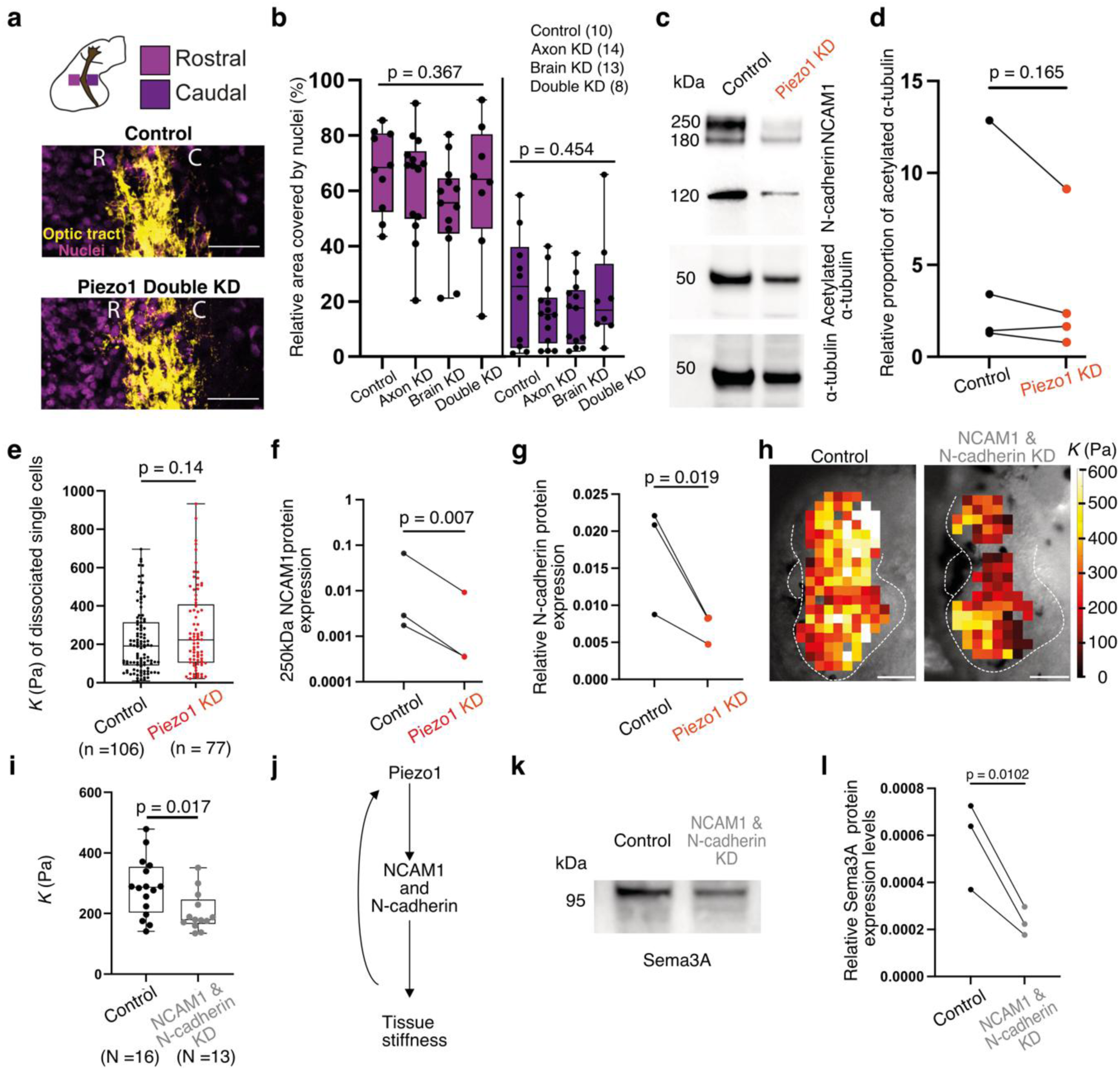
Piezo1 modulates tissue stiffness by regulating cell-cell adhesion. a-b, Piezo1 downregulation does not alter local cell body densities. **a,** Representative images of nuclei (DAPI, magenta) in brain tissue rostral (R) and caudal (C) to the optic tract (DiI, yellow) in stage 40 *Xenopus* embryos. Scale bars: 50 μm. **b**, The relative area occupied by nuclei rostral and caudal to the optic tract did not change after Piezo1 downregulation (two independent experiments; one-way ANOVA). **c,** Western blot image of NCAM1, N-cadherin, acetylated α-tubulin and α-tubulin protein expression in control and Piezo1-depleted brains. **d,** Western blot quantification showed that the relative proportion of acetylated α tubulin (acetylated α-tubulin/total α-tubulin) remained unchanged after Piezo1 knockdown. **e**, Quantification of the reduced apparent elastic modulus *K* of dissociated brain cells. Cell stiffness did not change upon Piezo1 downregulation (Mann-Whitney test). Each point corresponds to a single cell. **f-g**, Western blot quantification (normalised to total protein levels) of the levels of **f,** NCAM1 (250KDa) and **g,** N-cadherin. Both cell-cell adhesion proteins were significantly reduced in Piezo1-depleted brains. **h**, AFM-based stiffness maps overlaid on brightfield images of the brain for control and NCAM1 & N-cadherin-depleted brains. Scale bars: 100 µm. **i**, Decreasing NCAM1 and N-cadherin levels significantly reduced brain tissue stiffness (nested t-test). Each point represents the median *K* of an embryo. **j**, Schematic of the mechanism linking Piezo1 to tissue stiffness. Piezo1 regulates the protein levels of major cell-cell adhesion proteins such as N-cadherin and NCAM1, which in turn regulate tissue stiffness. **k**, Western blot image of Sema3A protein expression in control and NCAM1&N-cadherin-depleted brains. **l**, Quantification (normalised to total protein levels) revealed that downregulating NCAM1 and N-cadherin significantly reduced Sema3A protein expression levels. **d, f, g, l,** Data were compared using ratio paired *t*-tests. *p* values are indicated in the graphs. **b, e, i,** Boxplots show the 25^th^, 50^th^ (median), and 75^th^ percentiles; whiskers the spread of the data. N denotes the number of animals, n denotes the number of measurements. Brain KD: brain knockdown tissue, Double KD: double knockdown (optic tract and brain tissue depleted of Piezo1), KD: knockdown.

Since cell densities did not change, we next assessed whether the mechanical properties of cells within the tissue were altered after Piezo1 knockdown. As cytoskeletal components are key contributors to cellular stiffness^37^, we first used Western Blots to analyse cytoskeletal protein expression in control and Piezo1-downregulated brains. The expression of β-actin was unaffected by Piezo1 downregulation (*p =* 0.410, ratio paired t-test) (Fig. 2k and Supplementary Fig. 3b), while α-tubulin levels decreased following Piezo1 downregulation (*p =* 0.045, ratio paired t-test) (Fig. 4c and Supplementary Fig. 5a). However, recent studies showed that the levels of microtubule acetylation rather than absolute α-tubulin levels affect cell mechanics^38,39^. In *Xenopus* brains, the ratio of acetylated to total α-tubulin was unchanged between Piezo1-downregulated and control brains (*p =* 0.165, ratio paired t-test) (Fig. 4d and Supplementary Fig. 5b), suggesting that the mechanical state of cells was not affected by Piezo1-knockdown.

To confirm if tissue softening was a consequence of cell softening, we measured the stiffness of acutely dissociated neuroepithelial cells from the developing brain using AFM. Single cell stiffness was not altered upon downregulating Piezo1 (*p =* 0.14, Mann Whitney test) (Fig. 4e), indicating that cell-intrinsic mechanics indeed did not contribute significantly to the decrease in brain tissue stiffness observed upon Piezo1 downregulation.

Having ruled out changes in cell densities and cellular stiffness as the cause of tissue softening due to Piezo1 depletion, we hypothesized that reduced cell-cell interactions might contribute to the observed softening. Western blot analysis was used to quantify protein expression levels of the two major cell-cell adhesion molecules found in the developing neuroepithelium: neural cell adhesion molecule 1 (NCAM1)^40^ and N-cadherin^41^. While the 180 kDa band of NCAM1, likely representing the unmodified protein^42^, showed no significant change (*p =* 0.199, ratio paired t-test) (Fig. 4c and supplementary Fig. 5c), the 250 kDa polysialylated NCAM1^42^ band significantly decreased following Piezo1 depletion (*p =* 0.007, ratio paired t-test) (Fig. 4c, f). Additionally, N-cadherin expression significantly decreased following Piezo1 downregulation (*p =* 0.019) (Fig. 4c, g), indicating that Piezo1-mediated changes in cell-cell adhesion may contribute to the measured decrease in tissue stiffness (Fig. 3).

To substantiate this hypothesis, we downregulated both NCAM1 and N-cadherin by injecting translation-blocking morpholinos into both dorsal blastomeres of 4-cell stage-embryos. Western blot analysis confirmed reduction in both NCAM1 (*p =* 0.012, ratio paired t-test) and N-cadherin protein expression (*p =* 0.001) (Supplementary Fig. 5d-f), but not of the α-tubulin (*p =* 0.157) or β-actin (*p =* 0.227) loading controls (Supplementary Fig. 5d, g, h). The downregulation of NCAM1 and N-cadherin led to a significant decrease in tissue stiffness (median *K*control = 270 Pa, median *K*KD = 184 Pa; *p =* 0.017, Nested t-test) (Fig. 4 h, i). Collectively, our findings thus provide strong evidence for Piezo1 modulating tissue stiffness not via changes in cell-intrinsic mechanics or cell density, but rather via regulating cell-cell adhesions (Fig. 4j).

Moreover, downregulating NCAM1 and N-cadherin resulted in a significant decrease in Sema3A protein expression (*p* = 0.010, ratio paired t-test) (Fig. 4k, l), indicating that tissue softening is indeed sufficient to decrease the expression of long-range chemical guidance cues.

### Environmental stiffness regulates chemical guidance cue expression ex vivo

To test if stiffness could modulate chemical guidance cue expression independently of the decrease in Piezo1 expression, we exposed *ex vivo* brain tissue to different mechanical environments (Fig. 5a). We first quantified mechanical interactions between the tissue and its environment using 3D traction force microscopy^43^ (Fig. 5a, b). Tissue explants of wild type stage 37/38 embryos - when axons already respond to Sema3A^18^ and Slit1 - were embedded in soft (40 Pa) and stiff (450 Pa) 3D hydrogels (Fig. 5a, b)^43^ and matrix deformations quantified (see Methods).

**Figure 5:**
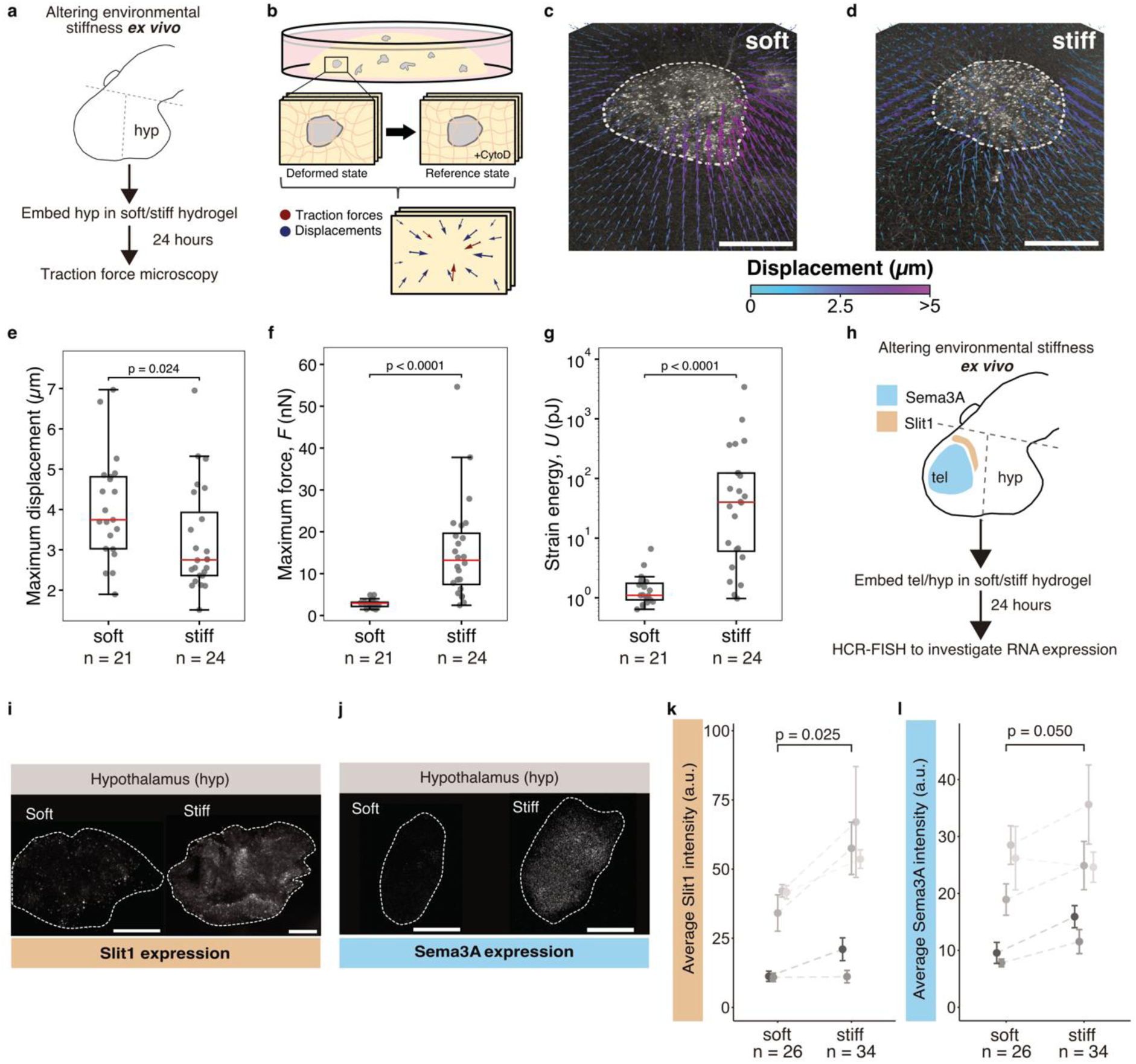
Altering environmental stiffness *ex vivo* affects traction forces and chemical guidance cue expression. a-b, Schematic of the experimental design. **a,** Hypothalamic brain tissue explants were dissected from *Xenopus* brains and embedded in collagen hydrogels for 24 hours. **b,** The matrix was imaged in a deformed and a reference state, the latter being induced by tissue relaxation using Cytochalasin D (CytoD) treatment, deformations quantified and forces calculated. **c-d,** Representative displacement fields for explants embedded in **c,** soft and **d,** stiff collagen hydrogels. Displacements are colour-coded, white dashed lines indicate tissue boundaries. Scale bars: 100 µm. **e,** The maximum matrix displacement was larger in soft than in stiff hydrogels. **f,** Explants locally exerted significantly larger maximum traction forces *F* on stiffer matrices compared to softer ones. **g,** The strain energy *U*, which is a measure of the work the tissue as a whole committed to deform the substrate, was significantly higher in stiff compared to soft hydrogels. For all graphs in **e-g**, each point represent single explants. Boxes correspond to the first and third quartile with a line at the median. Whiskers extend from the box to the farthest point lying within 1.5x the inter-quartile range. Data were compared using a Mann-Whitney U test. *p* values are indicated in the figure. **h,** Schematic of the experimental design for perturbing environmental stiffness *ex vivo*. The telencephalon and hypothalamus regions, were dissected (boundaries indicated in dashed lines) and embedded in soft or stiff 3D substrates. HCR-FISH was used to quantify Slit1 and Sema3A mRNA expression after 24 hours. **i-j,** Representative images of Slit1 (**i**) and Sema3A (**j**) expression in hypothalamic tissue embedded in soft (left) and stiff (right) substrates. Scale bars: 75 µm. **k-l,** Quantification of mRNA expression. Both Slit1 (**k**) and (**l**) Sema3A expression increased significantly in stiffer environments. Points represent means with standard errors of biological replicates. Ratio paired t-tests were used to compare groups. *n* denotes the number of tissue explants from 5 independent experiments.

Cells adhered to the surrounding matrix, dynamically pulled on it and visibly deformed the matrix (Fig. 5c-e, Supplementary movie 1), thus probing its mechanical properties. 3D traction force microscopy revealed that the tissue exerted significantly larger forces *F* on the stiff hydrogels than on the soft ones (median maximum *F*soft = 3 nN, *F*stiff = 13 nN; *p =* 2E-7, Mann-Whitney U test) (Fig. 5f and Supplementary movie 2). These forces were likely exerted collectively by multiple cells in the explant^44^. In line with larger local forces in the stiff matrices, the total work done by the tissue to deform the matrix was also significantly enhanced in stiff environments (median strain energies *U*soft = 1 pJ, *U*stiff = 40 pJ; *p* = 1E-6, Mann-Whitney U test) (Fig. 5g). As the forces exerted by cells are experienced by their own intrinsic force sensors, such as Piezo1, the observed increase in traction forces in stiffer environments suggested an enhanced activation of mechanosensing proteins, potentially regulating the expression of Sema3A and Slit1.

Thus, we next measured the effect of the environmental stiffness on Slit1 and Sema3A mRNA expression using HCR-FISH (Fig. 5h-l). Here, we used both stiff brain regions^19^ from the telencephalon and adjacent soft brain regions^19^ from the hypothalamus (Fig. 4a). After 24 hours in culture, Slit1 and Sema3A expression in the stiffer telencephalic tissue - which typically produces the guidance cues *in vivo* - was unaffected by environmental stiffness (*p*Slit1 = 0.824, *p*Sema3A = 0.542, ratio paired t-test) (Supplementary Fig. 6). However, in the softer hypothalamic tissue -which does not produce either guidance cue *in vivo*- we observed a significant increase in both Slit1 and Sema3A mRNA expression in stiff compared to soft substrates (*p*Slit1 = 0.025, *p*Sema3A = 0.050, ratio paired t-test) (Fig. 5i-l). These data showed that altering environmental mechanics is sufficient to modulate the expression of chemical guidance cues in soft brain tissues *ex vivo*.

### Environmental stiffness regulates Sema3A expression in vivo

To corroborate the effect of tissue stiffness on the expression of long-range chemical signals *in vivo*, we first employed a biochemical approach to increase the stiffness of brain tissue. We hypothesized that the application of lysophosphatidic acid (LPA), which enhances RhoA-mediated actomyosin-based contractility in neurons^48^, would increase tissue stiffness. Indeed, treatment of Stage 33/34 embryos with 100 µM LPA for 2-3 hours led to a significant increase in brain tissue stiffness (median *K*control = 216 Pa, median *K*LPA = 304 Pa; *p =* 0.01, Nested t-test) (Supplementary Fig. 7a-c). Consistent with the important role of tissue stiffness in regulating long-range chemical signal expression, Sema3A expression was significantly increased after 6 hours of this treatment (*p =* 0.05, unpaired t-test with Welch’s correction) (Supplementary Fig. 7d-f).

To verify *in vivo* that Sema3A mRNA expression was indeed induced by the stiffening of the tissue and not due to some biochemical side effect from the LPA treatment, we then exploited the fact that the elastic modulus of brain tissue increases under uniaxial compression, thus effectively resulting in a stiffer environment^45^. To locally stiffen soft hypothalamic brain regions of stage 35/36 embryos, we used AFM to apply compressive forces of 30 nN for > 6 hours (until embryos reached stage 40) *in vivo*^19,46^.

Control measurements, in which a force of 10 nN was applied to embryonic brains with no prior compression (Supplementary Fig. 8a(*i*), b) and to the same brains after an initial indentation with a force of 30 nN for 15 minutes (mimicking the strain stiffening conditions) (Supplementary Fig. 8a(*ii*), c), confirmed that the stiffness *k* of the tissue immediately increased under compression (*p* = 0.001, Repeated measures ANOVA followed by Tukey’s multiple comparison) (Supplementary Fig. 8d-e), likely because of the non-linear elasticity of the tissue and water flow. Furthermore, using a standard linear model fit (see Methods) to the indentation-time curves obtained above (Supplementary Fig. 8f), we found an increase in the tissue’s apparent viscosity *η* under compression (*p* < 0.001, repeated measures ANOVA followed by Tukey’s multiple comparison) (Supplementary Fig. 8g-h), likely because of fluid exudation (loss of interstitial fluid), which increases internal friction and resistance to further deformation^47^. However, changes in the mechanical properties of tissue under compression were transient, as the tissue’s apparent elastic moduli *K* were similar before and after compressing the brains for >6 hours (*p* = 0.499, one sample t-test) (Supplementary Fig. 8i).

We then assessed mRNA expression in the compression-stiffened brains using HCR-FISH (Fig. 6a). Slit1 expression was not significantly different from controls in compression-stiffened brains (*p =* 0.169, unpaired t-test with Welch’s correction) (Supplementary Fig. 9). However, in line with our *ex vivo* data (Fig. 5l), stiffening the tissue resulted in the ectopic production of the chemical guidance cue Sema3A at the hypothalamus (Fig. 6c, d). In compression-stiffened brains, there was a significant increase in mRNA levels compared to controls (*p =* 0.011, unpaired t-test with Welch’s correction) (Fig. 6b-d), indicating that *in vivo* tissue stiffness may indeed regulate the expression of the long-range chemical signal, Sema3A.

**Figure 6:**
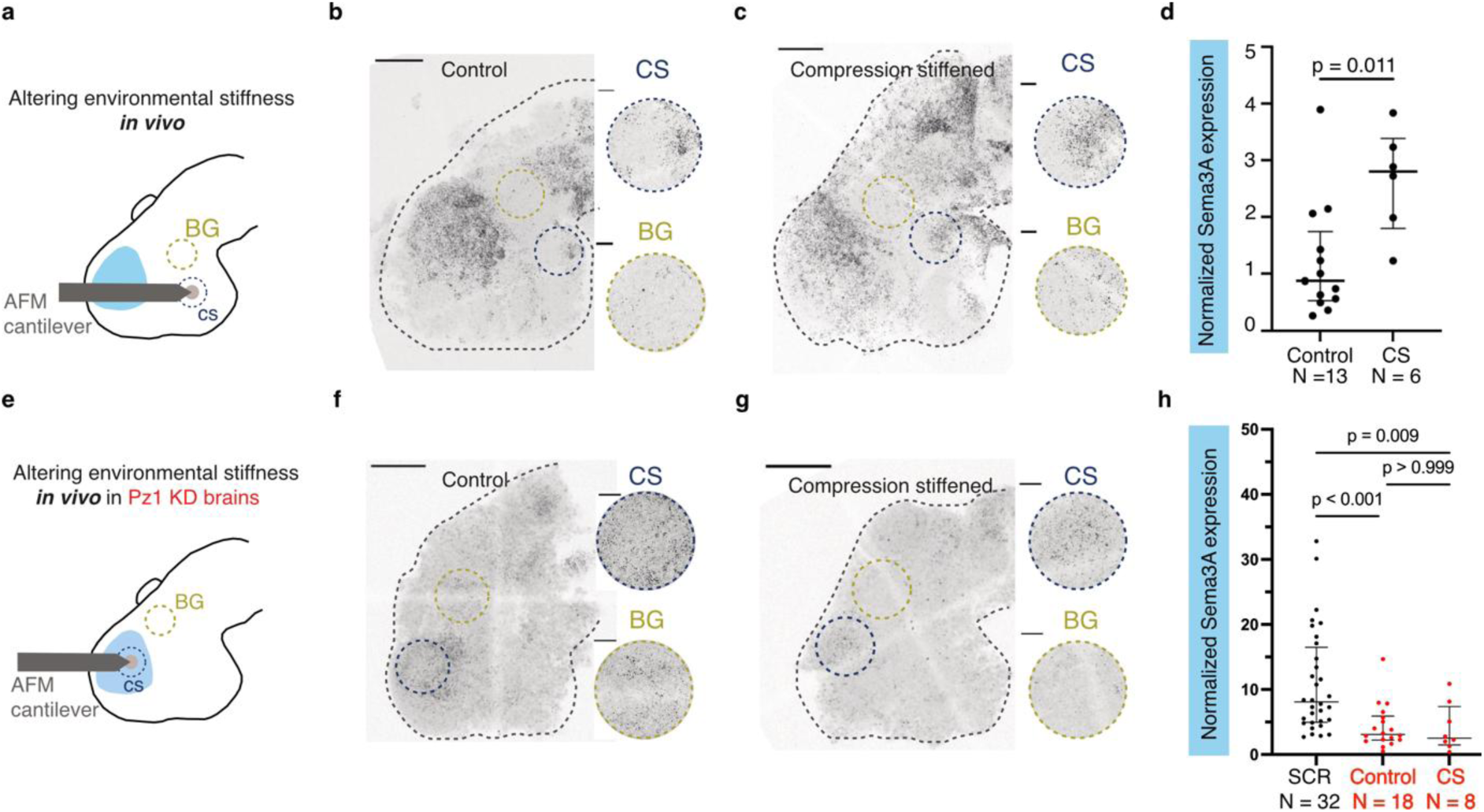
Altering environmental stiffness *in vivo* affects Sema3A expression. **a,** Schematic of the experimental setup for locally increasing tissue stiffness in wild type brains *in vivo.* The hypothalamus region of the brain was compression-stiffened with a sustained force of 30-40 nN applied with an AFM probe of ∼90-120 µm diameter for > 6 hours. **b-c,** Representative images of Sema3A HCR-FISH in control and compression-stiffened brains. The insets show representative regions which were selected for analysis. **d,** The ratio of the total area covered by signal in the compression-stiffened (CS) region to the mean area covered by the signal in background (BG) regions were analysed. Sema3A expression significantly increased in compression-stiffened hypothalamic brain regions compared with control brains. An unpaired t-test with Welch’s correction was used for statistical analysis, the resulting p value is indicated in the figure. **e,** Schematic of the experimental setup for perturbing tissue stiffness in Piezo1 knockdown brains *in vivo*. The telencephalic region of the brain was compression-stiffened as in (**a**). **f-g,** Representative images of Sema3A HCR-FISH in control and compression-stiffened Piezo1 knockdown brains. The insets show representative regions which were selected for analysis. **h,** Sema3A expression decreased in the telencephalon of Piezo1 KD-brains; expression could not be rescued by compression-stiffening the tissue, suggesting that Piezo1 is not only involved in regulating tissue stiffness but it is also required to sense and respond to increased tissue stiffness. Kruskal-Wallis test, p < 0.0001, followed by Dunn’s post-hoc test for multiple comparison (p-values provided in the figure). Each point refers to an embryo. Scale bars in whole brain images: 100 μm; scale bars in insets: 20 μm. hyp: hypothalamus, KD: knockdown, tel: telencephalon.

Since Piezo1-depleted brains had decreased Sema3A expression (Fig. 2) and were softer than control brains (Fig. 3), and as an increase in tissue stiffness led to an increase in Sema3A expression *in vivo* (Fig. 5j, l), we finally tested if stiffening telencephalic regions, where Sema3A is normally produced, is sufficient to rescue Sema3A expression in Piezo1 knockdown brains (Fig. 6e). Compression-stiffening Piezo1-depleted stage 35/36 brains for > 6 hours did not restore Sema3A levels (*p*Control vs CS > 0.999; Kruskal-Wallis test followed by Dunn’s post-hoc test) (Fig 6f-h), suggesting that both tissue stiffness and adequate levels of Piezo1—which not only regulates but also detects tissue stiffness—are required to lay down the appropriate chemical landscape for supporting axon pathfinding in the developing brain.

## Discussion

During numerous processes in diverse biological systems, including brain development, cells encounter a plethora of chemical and mechanical cues in their environment, which they detect, integrate, and interpret. We here discovered an intricate interplay between mechanical and long-range chemical signalling *in vivo*, with both influencing each other, showing that neither cue can be fully understood in isolation. Importantly, local tissue stiffness regulates the availability of diffusive, long-range chemical signals, thereby impacting cell function at sites distant from the origin of mechanical signalling.

In tissues, cells constantly and dynamically exert forces on their environment. These contractile forces increase with increasing tissue stiffness (Fig. 5f). Larger forces, in turn, induce more augmented deformation of the cells’ intrinsic force sensors, such as Piezo1, thereby leading to their activation^49^. We found that Piezo1 serves at least two functions in the developing brain. First, it is involved in sensing tissue stiffness in both RGC axons^19^ (Fig. 1d,h) and the surrounding neuroepithelial cells (Figs. 1e, 6h). Second, it inherently contributes to regulating the mechanical properties of the tissue (Fig. 3a-d). The appropriate tissue stiffness in turn triggers the expression of long-range chemical guidance cues (Figs. 5, 6).

Downregulation of Piezo1 in brain and skin tissues led to a decrease in tissue stiffness (Fig. 3 and Supplementary Fig. 4) and a subsequent decrease in the expression of Sema3A and Slit1 in the brain (Fig. 2). In contrast, the downregulation of Sema3A was followed by an increase in tissue stiffness (Fig. 3e-h), ruling out that the observed tissue softening following Piezo1 depletion was the direct consequence of decreased Sema3A expression.

The reason for the tissue stiffening following Sema3A depletion is currently unknown. Sema3A is known to decrease cell-ECM adhesion in various cell types, including neural crest cells^50^. Furthermore, Sema3A has been shown to decrease cell-cell adhesion via the internalisation of cadherins^51^. Since our data show that cell junctions are an important regulator of tissue stiffness (Fig. 4), and Sema3A destabilises cell junctions, Sema3A depletion might lead to the stabilisation of cell junctions and thus to brain tissue stiffening. Regardless, our data show that feedback mechanisms between tissue mechanics and gene expression levels do not always act reciprocally.

Recent work has shown that environmental stiffness may regulate expression levels of Piezo1. For example, softer substrates lead to a decrease in Piezo1 expression levels *in vitro*^52,53^, Piezo1 levels in microglial cells correlate with the stiffness of Aβ plaque-associated brain tissues^54^, and in macrophages Piezo1 levels scale with the stiffness of ischemic tissues^55^. We here found the opposite effect: a regulation of environmental stiffness by Piezo1 expression levels. A decrease in Piezo1 expression led to tissue softening in a developing vertebrate system *in vivo* (Fig. 3 and Supplementary Fig. 4), which was previously only observed in genetically induced *Drosophila* gliomas but not in non-transformed *Drosophila* brains^56^.

While Piezo1 has been shown to regulate cell proliferation^57,58^ in other *in vivo* systems and the mechanical properties of epithelial cells^59^ *in vitro*, this was not observed here.

Piezo1 depletion in the developing *Xenopus* brain *in vivo* had no effect on local cell densities or cell stiffness (Fig. 4 a-e). Instead, Piezo1 downregulation resulted in a significant depletion of cell-cell adhesion (Fig. 4c, f-g), accompanied by a decrease in tissue stiffness and lower Sema3A levels (Figs. 2, 3, 4h-l) – consistent with an important role of tissue stiffness in regulating the expression of long-range chemical signals. The simultaneous downregulation of Piezo1 and upregulation of NCAM1 and N-cadherin to rescue the phenotype in developing *Xenopus laevis* brains was not feasible, however the downregulation of these cell-cell adhesion molecules was sufficient to reduce tissue stiffness (Fig. 4h-i) and subsequently Sema3A expression (Fig. 4k-l), indicating that the mechanoregulation of tissue stiffness is not specific to Piezo1 but potentially mediated by NCAM1 and N-cadherin.

However, Piezo1 was required to induce Sema3A expression in response to tissue stiffness exceeding a critical threshold (Fig. 6h). While increasing tissue stiffness *in vivo* biochemically (Fig. S7) or mechanically (Fig. 6) was sufficient to induce ectopic Sema3A expression in soft brain regions of wild type embryos, tissue stiffening in Piezo-downregulated brains tissue failed to rescue Sema3A expression (Fig. 6), indicating that Piezo1 is required by the brain parenchyma to detect mechanical signals and induce Sema3A expression (Fig. 6h).

How Piezo1 activity contributes to regulating Sema3A expression remains to be studied. As Piezo1 is a non-selective cation channel that gives rise to calcium transients^26^, a regulation via calcium-dependent transcription factors such as activator protein (AP)-1^60^ or calcium-dependent enzymes such as mitogen-activated protein kinase (MAPK)/extracellular signal–regulated kinase (ERK) kinase (MEK) 1/2^60^, which are involved in the regulation of Sema3A expression^60^, is conceivable. Furthermore, softening tissue by depletion of Piezo1 or cell-cell adhesion proteins early in development may not only affect Piezo1-mediated mechano-responses but potentially also neural stem/progenitor cell fate determination^25,61^ or FGF signalling^62^, which regulates the expression of both Sema3A and Slit1^17,35^. Future work will reveal the precise molecular details on how tissue mechanics and long-range chemical signalling impact each other.

A dependence of Sema3A expression on tissue stiffness is consistent with expression patterns found in healthy *Xenopus* brains. Normally, Sema3A is expressed in the telencephalon, which is significantly stiffer than the diencephalon and hypothalamus^19^, where Sema3A is not expressed (Fig. 2). Yet, Sema3A is already expressed in the telencephalon region at stage 32^35^ - before the telencephalon stiffens^20^ – indicating that other factors contribute to Sema3A regulation. *Ex vivo*, environmental stiffness affected Sema3A and Slit1 mRNA expression in the soft hypothalamus, which does not produce these guidance cues, but had no effect on the stiffer, guidance cue-producing telencephalic brain regions (Supplementary Fig. 6). These data suggested a sigmoidal relationship between tissue stiffness and the mechanical induction of guidance cue expression. Namely, at lower stiffness, expression remains low. However, upon reaching a certain critical stiffness threshold, chemical guidance cue expression is induced. Beyond this threshold, further increases in environmental stiffness have no additional effect on Sema3A expression levels. Hence, once the tissue is sufficiently stiff to produce the cue, further increasing environmental stiffness will not affect its expression.

Collectively, our data show that in the developing *Xenopus* brain, Piezo1 is critical for setting up the mechanical and, consequently, chemical landscape required for proper RGC axon guidance; it is involved in both RGC-intrinsic and cell-extrinsic regulations of axon pathfinding in the developing retino-tectal system. Our findings raise the intriguing possibility that tissue mechanics may regulate the transcription of many other diffusible long-range biochemical factors^63^ in various other organ systems – and *vice versa*. Through such bidirectional regulation, tissue mechanics could thus not only directly (i.e. through mechanotransduction pathways) but also indirectly (i.e. through the regulation of chemical signalling pathways) impacting cell function and tissue development over large distances. Understanding the interplay between chemical and mechanical signalling is challenging but has great potential to provide new insights into the regulation of development, physiology, ageing, and disease.

## Methods

Detailed methods are available in the online version of the paper, including the associated code and references.

## Data availability

The datasets generated during and/or analysed during the current study are available from the authors on reasonable request.

## Acknowledgements

We thank Christine Holt, Maximilian Jakobs, Sarah Foster, Buzz Baum, Kevin Chalut, and Nick Brown for insightful discussions; Liz Williams, Sebastian Vasquez Sepulveda, Isaac Baguley, Asha Dwivedy, Chaitanya Dingare, Can Aztekin (EPFL Lausanne, Switzerland), Ben Fabry, David Böhringer, Johannes Rheinlander, Ryan Greenhalgh, and Jakub Sedzinski (The Danish Stem Cell Centre, University of Copenhagen, Denmark) for technical help and advice. We also acknowledge the Cambridge Advanced Imaging Centre for support in this work. This work has been funded by a Wellcome Trust PhD studentship 222280/Z/20/Z (S.M.), Cambridge Trust scholarship (S.M.), an MRC Doctoral Training Grant awarded to the Wellcome-MRC Cambridge Stem Cell Institute: MR/R50211X/1 (R.J.M.), the European Research Council Consolidator Award 772426 MECHEMGUI (K.F.), the European Research Council Synergy Grant 101118729 UNFOLD (K.F.), the German Research Foundation (DFG) projects 460333672 CRC1540 EBM and 270949263 GRK2162 (K.F.) and an Alexander von Humboldt Professorship (Alexander von Humboldt Foundation) (K.F.).

## Author contributions

E.K.P., S.M., and K.F., designed the study. E.K.P., S.M., N.G., R.J.M., K.A.M., J.M.B, A.W, and A.J.T., carried out experiments, data analysis, data curation and visualization. E.K.P, S.M., and K.F., wrote the manuscript. K.F., supervised the study. All authors reviewed and edited the manuscript.

## Authors’ notes

The first two authors should be regarded as joint co-first authors. Co-first authors can prioritize their names when adding the paper’s reference to their CVs.

