## Supplementary figures and images for "Long-range chemical signalling *in vivo* is regulated by mechanical signals"

### Movie S1

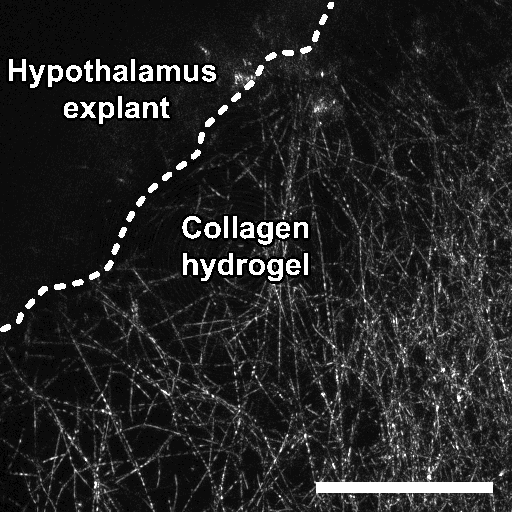

### Movie S2

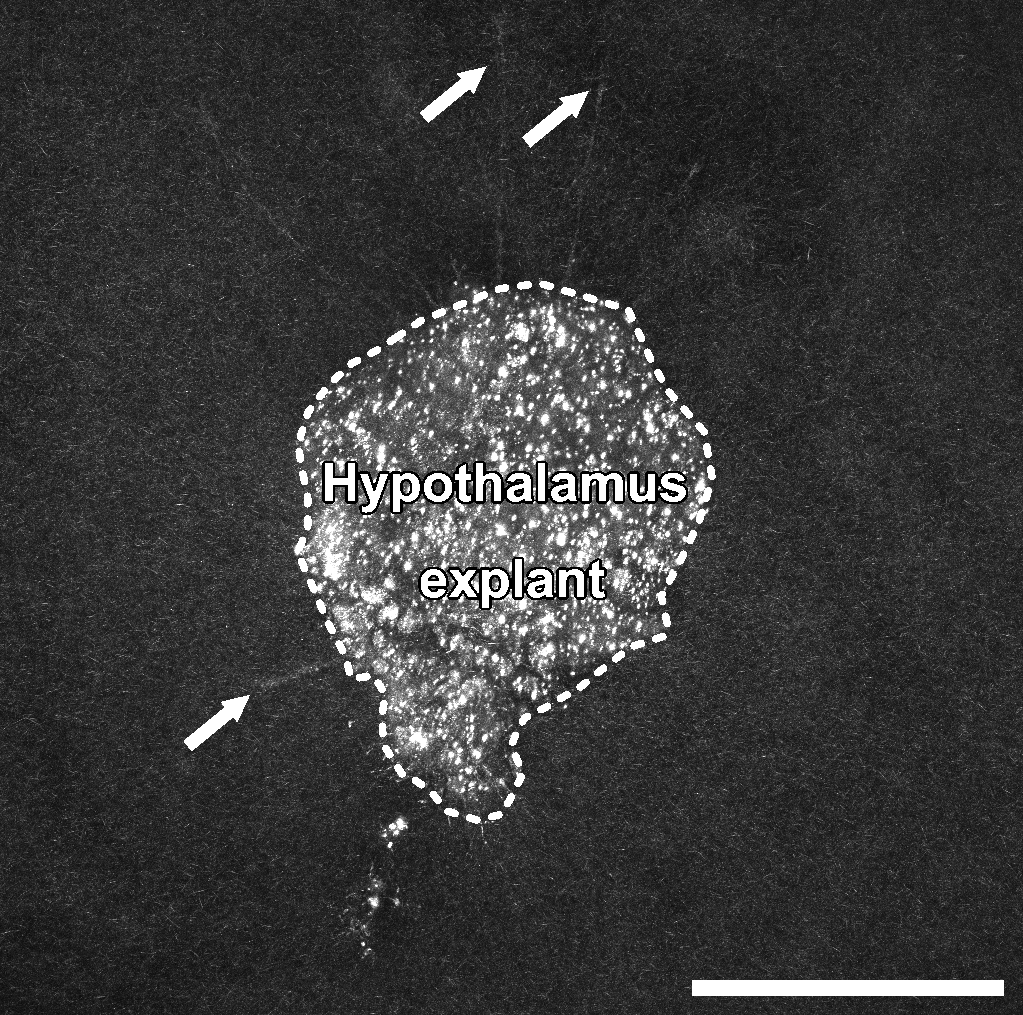
