## Supplement for "Long-range chemical signalling *in vivo* is regulated by mechanical signals"

### Online Methods

All reagents were obtained from Sigma-Aldrich unless otherwise specified.

#### *Animal experiments*

##### *Animal model*

All animal experiments were approved by the Ethical Review Committee of the University of Cambridge, in compliance with guidelines set by the UK Home Office. Wild-type *Xenopus laevis* embryos of both sexes were obtained via *in vitro* fertilisation. Embryos were reared in 0.1 × Marc's Modified Ringer Solution (MMR) or 0.1 × Modified Barth's Saline (MBS) at 14–18°C to reach the desired developmental stage, according to Nieuwkoop and Faber staging[1]. All animals used in this study were below embryonic stage 45.

##### *Morpholino injection*

Freshly fertilised embryos were chemically dejellied in 2% w/v Cysteine in 0.1 × MMR/MBS, pH 8.0, for 2-5 minutes. Embryos were visually inspected under a stereomicroscope and when the jelly coat was no longer visible, embryos were rinsed in several washes with 0.1 × MMR/MBS. The dejellied embryos were gently transferred to a mesh-bottomed dish containing 4% w/v Ficoll in 0.1 × MMR/MBS, pH 7.5. Glass capillaries (1.0 mm OD by 0.5 mm ID, Harvard Apparatus) were pulled into needles and loaded with the morpholino construct. Needles were then affixed to a micromanipulator (connected to a FemtoJet 4X microinjector (Eppendorf)) and manually broken with forceps. The droplet size was calibrated in mineral oil, using a reticule in the microscope eyepiece, to dispense ~5 nL of morpholino. Fluorescein or lissamine-tagged translation-blocking morpholino oligonucleotides (MOs) targeting Piezo1 Tropicalis (5′-CACAGAGGACTTGCAGTTCCATCCC-3′)[2], Piezo1.L- Laevis (5'- CGCACAGGACTTGCAGTTCCATCCC -3')[3], NCAM1 (5′-GATCCTTAATGTGCAGCATTGTAA-3′), and N-cadherin (5′- GAAGGGCTCTTTCCGGCACATGGTG-3′) were designed and synthesized by GeneTools (GeneTools, OR, USA). 15 ng of either Piezo1 MO, NCAM1 or Control MO (GeneTools, OR, USA; 5′-CCTCTTACCTCAGTTACAATTTATA-3′) and 7.5 ng of N-cadherin MO was injected into the dorsal blastomeres of 4-cell-stage embryos, as previously described[4]. For whole nervous system downregulation, both dorsal blastomeres were injected; for tissue-specific downregulation, one of the two dorsal blastomeres was injected. Embryos were reared until the requisite stage for use and, prior to downstream experiments, screened using a fluorescence stereomicroscope. Only embryos presenting a clear signal in the tissues of interest were selected.

***Exposed brain preparation***

Embryos were anaesthetised in *Xenopus* exposed brain medium (0.04% w/v tricaine methanesulfonate (MS222), 1 × PSF in 1.3 × MBS/MMR, pH 7.6)[5], this higher osmolarity retards skin growth. Embryos were transferred to a Sylgard coated Petri dish and immobilized laterally with bent 0.2 mm minutien pins (Austerlitz). Skin, dura, and eye were dissected out using fine forceps and 0.1/0.15 mm minutien pins (Austerlitz) in pinholders, to expose the brain from the dorsal to ventral midline and from the hindbrain to the telencephalon. Embryo viability throughout all *in vivo* experiments was assessed by the presence of a visible heartbeat, prior to processing the samples for downstream analysis.

***Lysophosphatidic acid treatment***

Stage 33/34 embryos were anaesthetised, and their brains were exposed as above and incubated for (*i*) 2-3 hours for AFM-based stiffness mapping, and (*ii*) 6 hours for HCR-FISH, in exposed brain medium (0.04% w/v tricaine methanesulfonate (MS222), 1 × PSF in 1.3 × MBS/MMR, pH 7.6)[5] containing either 100 µM DMSO (control) or 100 µM Lysophosphatidic acid (LPS; 1-oleoyl-2-hydroxy-*sn*-glycero-3-phosphate; Avanti Research). Embryo viability was assessed by confirming the presence of a visible heartbeat and the samples were then processed for downstream analysis.

***Brain-targeted electroporation***

Finely controlled spatio-temporal knockdown of Semaphorin3A (Sema3A) was achieved using a previously described targeted electroporation method[6]. The electroporation, rather than blastomere injection, approach was used for better spatio-temporal control of the downregulation and to avoid other developmental defects related to early Sema3A depletion[7]. Vitelline membranes were manually removed from stage 28 embryos using fine tipped forceps. Stage 29/30 was chosen as axons had yet to grow across the contralateral brain surface, ensuring sufficient Sema3A depletion by the time the axons would encounter the cue.

Embryos were anaesthetised in (0.04% w/v tricaine methanesulfonate (MS222), 1 × Penicillin- Streptomycin-Amphotericin B (PSF, Lonza, 1% v/v of a 100 x solution containing: penicillin (10,000 U/mL), streptomycin (10,000µg/mL), amphotericin B (25µg/mL)) in 1.0 × MBS, pH 7.6), and an individual embryo was placed ventral side down in a custom fabricated, Sylgard electroporation chamber with 0.5 mm wide platinum electrodes on either side. Glass capillaries (1.0 mm OD by 0.5 mm ID) were loaded with 1 mM fluorescein-labelled validated Sema3A MO[8] (GeneTools,OR, USA; 5'- TGCAATCCAGGTCAGAGAGCCCATG

-3') or standard control morpholino solution (GeneTools, OR, USA; 5’-CCTCTTACCTCAGTTACAATTTATA-3’) and placed in a micromanipulator connected to a picospritzer (Intracel). The fluorescein label of the constructs enabled the visualization of their localization.

The capillary was inserted between the eye and brain, such that the target region was between the positive electrode and capillary. Eight pulses of morpholino were dispensed, followed by eight 18 V shocks, for a duration of 50 milliseconds, spaced one second apart using a TSS20 Ovodyne electroporator (Intracel). The electrodes were then removed, and the embryo was gently transferred with a Pasteur pipette to a dish containing 0.1 × MBS and left to develop at 20°C to stage 40. Embryonic brains were exposed (see Exposed brain preparation) and screened for fluorescence at the telencephalon region of the brain. Embryos with signal at the region of the brain where Sema3A is normally expressed were selected for AFM measurements (Fig 3e).

***Xenopus laevis whole brain immunohistochemistry***

4-cell stage embryos were injected with Piezo1 MO in one of the two dorsal blastomeres. Stage 40 embryos were screened; the fluorescent side corresponded with MO-injected (and therefore Piezo1 depletion), whereas the non-fluorescent side had wild type levels of Piezo1. Embryos were fixed in 4% PFA in 1 × PBS for 4 hours at room temperature, followed by two washes in 1 × PBS. The brains were then dissected out in PBST solution (1 × PBS with 0.1% TritonX-100), using 0.1 - 0.15 mm minutien pins (Austerlitz). The dissected brain samples were washed in PBST solution containing 0.2% BSA (bovine serum albumin and blocked for 45 minutes at room temperature in blocking solution (PBST solution containing 0.2% BSA and 10% donkey serum). Samples were then incubated with rabbit anti-Piezo1 antibody (Proteintech, 28511-1-AP, 1:10 dilution in blocking solution) at 4°C overnight. Samples were washed thrice in PBST solution for 30 minutes before adding the secondary antibody, anti-rabbit Alexa Fluor 594 (Abcam, ab150080, 1:300 dilution in blocking solution) for 2 hours at room temperature. Samples were washed thrice in PBS and incubated with DAPI (1:10,000) in 1 × PBS for 15 minutes. The samples were then washed five times in 1 × PBS. Following this, the samples were upgraded serially, with 15 minute washes, in 30, 50 and 70 % glycerol. Samples were mounted, in 35 mm glass bottomed dishes (Ibidi), laterally/dorsally as required. Both MO-injected and non-injected sides of the embryos were imaged (Leica SP8 confocal microscope; 40x oil, NA =1.3; z-step size = 2 µm). Maximum intensity projections of the first 40 µm of each z-stacks was made in Fiji. The telencephalon was identified using the DAPI channel; the intensity of Piezo1 expression in the telencephalon was measured. Statistical analysis was done with GraphPad PRISM.

#### *Visualising RGC axons*

##### *RGC axon labelling in vivo*

Stage 40 embryos were fixed using 4% paraformaldehyde (PFA; Thermo Fisher Scientific) in 1 × phosphate buffered solution (PBS; Thermo Fisher Scientific) for 2 hours at room temperature or overnight at 4°C. Post-fixation, embryos were rinsed in 1 × PBS and immobilized with bent 0.2 mm minutien pins (Austerlitz) on Sylgard coated dishes. DiI (1,1'-Dioctadecyl-3,3,3',3'- Tetramethylindocarbocyanine Perchlorate, Molecular Probes) crystals were dissolved in 100% ethanol and heated for a few minutes at 65°C. The DiI solution was then loaded into microinjection glass capillaries (1.0 mm OD by 0.78 mm ID, Harvard Apparatus) and connected to a FemtoJet 4X microinjector (Eppendorf). DiI was injected into the eye, between the boundary of the lens and retina, until a full roseate ring formed as described in[9]. After 24-28 hours in 1 × PBS at room temperature, the brains were dissected and mounted in 1 × PBS in the lateral view. Z-stack images were obtained using a confocal microscope (SP8, Leica Microsystems, UK; 20× air, NA = 0.75; z-step size = 1 µm).

***RGC axon phenotype analysis***

Only samples that were mounted with the brain positioned properly and with visible tracts were used in the phenotype and elongation analysis (below). DiI-labelled images were randomized and OTs with normal projections (axons reach the tectum, without mid-OT bend defects or visibly straying axons), or that displayed defects such as mid-OT bend stalling, or axon misprojections were counted. Misprojections frequently occurred near the mid-OT bend but also included axons straying in other parts of the tract. Some of the misprojections included deviating at the mid-OT bend, bypassing the bend, avoiding the tectum or projecting into the telencephalon. Percentages of each phenotype were compared for the various experimental conditions.

***OT elongation analysis***

Maximum projections were made across 10-15 µm confocal image stacks of wholemount brains with DiI-labelled optic tracts (OT). The OTs were manually outlined in Adobe Illustrator. Elongation of the OT was calculated by the major-to-minor axis ratio using a previously described[2] automated MATLAB algorithm. Briefly, the major and minor axes were determined by fitting ellipses, with the same normalized second central moment as the OT area, around the OTs.

#### *Visualising mRNA expression in situ*

***In situ hybridization***

Whole-mount in situ hybridization was adapted from[10]. Stage 40 embryos were collected in cold PBS, fixed with 4% PFA (Thermo Fisher Scientific) for 2 hours at room temperature. Whole brains were carefully dissected out in nuclease-free PBS (Thermo Fisher Scientific) and washed thrice with cold PTw (0.1% Tween-20 in RNase-free PBS). Samples were transferred to a 12- well plate with mesh adapters and treated with 20µg/ml of proteinase K for 5 minutes at room temperature, then washed twice for five minutes in freshly prepared 2 mg/ml glycine in PTw. Brains were post-fixed in 0.2% glutaraldehyde in PBS, for 20 minutes at room temperature. Samples were washed thrice for five minutes in PTw. After treatment with freshly prepared 0.1% sodium borohydride in PTw for 20 minutes, brains were washed a further three times in PTw, and twice in hybridization buffer (50% formamide (Thermo Fisher Scientific), 750 mM NaCl, 1 × PE (10mM PIPES pH6.8; 1mM EDTA), 100µg/ml yeast t-RNA (Thermo Fisher Scientific), 0.05% heparin and 1% SDS in nuclease-free water) for five minutes at room temperature. Samples were pre-hybridized for at least 1 hour at 63°C in a hybridization oven, in a humidity chamber, and samples were hybridized with 2µg/ml of DIG-labelled Sema3A probe in hybridization solution overnight at 63°C. A pCS2+-*XenopusSema3A*[8] plasmid, from the lab of Christine Holt, was used as a template to prepare the DIG-labelled RNA probe. Samples were then washed in hybridization buffer at 63°C for 30 minutes, and twice for 30 minutes with 300 mM NaCl, 1 × PE, and 1% SDS at 63°C. Followed by two 30-minute washes in 50 mM NaCl, 1 × PE, and 0.1% SDS at 50°C, and a brief rinse in NTE buffer (500 mM NaCl, 10 mM Tris-HCl pH 8.0; 1mM EDTA). The samples were then treated with 100µg/ml RNaseA (Thermo Fisher Scientific) and 100U/mL of RNaseT1 in NTE for 60 minutes at 37°C and rinsed briefly in NTE. The samples were then washed in (50% formamide, 300 mM NaCl, 1 × PE, 1% SDS) at 50°C for 30 minutes, followed by (50% formamide, 150 mM NaCl, 1 × PE, 0.1% Tween-20) at 50°C for 30 minutes. Finally, the samples were washed twice at room temperature, and once for 20 minutes with (500 mM NaCl, 1 × PE, 0.1% Tween-20) at 70°C to inactivate endogenous alkaline phosphatases. Samples were blocked in 1 × MABT (0.1 M maleic acid, 150 mM NaCl, 0.1% Tween-20, pH 7.5), with 2 mM levamisole (Abcam), 2% Boehringer blocking reagent (Roche), and subsequently incubated overnight at 4 °C with a 1:5,000 dilution of anti-digoxigenin antibody (Roche; 11093274910) in blocking reagent. After thorough washings, thrice quickly and 5-6 times for an hour each with freshly made 2mM levamisole in 1 × MABT, samples were washed twice for 20 minutes with 2mM levamisole in NTMT (100 mM NaCl, 100 mM Tris pH 9.5, 50 mM $MgCl_2$, 0.1% Tween-20). The colour reaction was initiated by removing the NTMT and adding BM Purple (Roche) to the brains and stopped with the changes of PTw containing 1 mM EDTA. Samples were mounted laterally in PBS and imaged using a Leica MZFLIII stereomicroscope with QImaging microPublished Color RTV camera, and QCapture Pro 7 software.

***Whole brain and in vitro tissue culture HCR-FISH***

To visualize RNA in fixed *Xenopus* tissue, a modified Hybridisation Chain reaction (HCR) RNA-FISH protocol[11] was employed. Embryos/embedded tissues were fixed in modified Carnoy's fixative (6 parts 100% ethanol, 3 parts 37% formaldehyde (Fisher chemical) and 1-part glacial acetic acid (Fisher chemical))[12] for 4 hours at room temperature or overnight at 4°C. Samples were rinsed in 70% ethanol in PBST (PBS + 0.1% Tween-20) for 10 minutes and dehydrated by three 30 minute washes with 100% ethanol. Thereafter, samples were stored at -20°C and then rehydrated serially with 70%, 50% and 25% ethanol in PBST for 15 minutes each. Samples were then rinsed in PBST thrice for 15 minutes. For embryos, samples were kept in PBST and brains were dissected out with 0.1/0.15 mm minutien pins (Austerlitz) in pinholders. For embedded tissues, the samples were dissected out of the 3D gel matrix using 0.1/0.15 mm minutien pins (Austerlitz) in pinholders. The dissected brains/tissues were transferred to a 4-well plate and bleached in a solution of 5% formamide, 2.5% 20 × SSC (Promega) and 28% $H_2O_2$ for 45 minutes under a Zeiss™ Cold Light Source at 100% intensity. The samples were gently rinsed and then washed thrice in PBST for 10 minutes each. They were then treated for 10 minutes at room temperature with 5 mg/ml proteinaseK  diluted in PBST, followed by three 5 minute washes with PBST. They were fixed again with a 20 minute wash with 3.7% formaldehyde in PBST and washed five times for 5 minutes each in PBST.

The samples were washed in 1 mL probe wash buffer (Molecular instruments: MI) (preheated to 37°C) for 5 minutes at room temperature and then washed in 500 mL hybridisation buffer for 30 minutes at 37°C. The probe solution was prepared by mixing custom probes (MI) in 500 mL of hybridisation buffer (MI). Final concentration was 24 nM for sema3A and 12 nM for slit1. The probe mixture was heated at 37°C for 30 minutes. The probe solution was then added to the samples and incubated overnight at 37°C. Samples were kept stationary. The following day, the wash buffer was preheated to 37°C in a water bath. Excess solution was removed, and the samples were washed twice for 30 minutes each in 1 ml of preheated wash buffer at room temperature. The samples were then washed for 5 minutes in 50% 5 × SSCT (5 x SSC (diluted in PBS) + 0.1% Tween 20) + 50% wash buffer at room temperature, following which they were washed with 5 × SSCT, two times 20 minutes each, at room temperature. Thereafter, they were pre amplified with 1ml amplification buffer (MI) for 10 minutes. 4 µl of H1 and 4 µl H2 hairpins, from 3 µM hairpin stocks (MI) were taken and separately heated at 95°C for 90 seconds and cooled for 30 minutes at room temperature, in dark. 8 µl of the prepared hairpin solution was mixed with 500 µl amplification buffer. The final concentration of the hairpins was 48 nM. Excess amplification buffer from the samples were removed and the hairpin mix was added and incubated at room temperature, overnight, in the dark. The samples were washed with 5 × SSCT, 2 times, 30 minutes each in 1 × PBS. They were then incubated with DAPI 1:10,000 solution in PBS, for 10 minutes and washed with 1 × PBS two times for 10 minutes each. The samples were then serially upgraded with 15 minute washes/until samples sunk to the bottom of the dish in 30%, 50% then 70% glycerol in PBS. Samples were stored at 4°C in 70% glycerol. The samples were mounted (laterally for whole brains) in 70% glycerol and 80 µm z-stacks were acquired on a laser scanning confocal microscope (SP8, Leica Microsystems, UK; 40× oil, NA = 1.3; z-step size = 2 µm). For experiments involving LPA treatment, brains were imaged on an inverted fluorescence light microscope (Thunder DMi8, Leica Microsystems, UK; 63× oil, NA = 1.4; z-step size = 1 µm). Multiple tiles were acquired for each sample, using the Leica LasX navigator software, and tile stitching was performed in the Leica LAS X software.

***HCR-FISH quantification***

To quantify Slit1 expression in whole brains, a sum intensity projection was obtained for 30 µm from the z stack where DAPI first appears. This sum intensity projection was used for further analysis. A rectangular ROI covering the Slit1 producing region at the boundary of the diencephalon and telencephalon was selected. The area covered by puncta was calculated using the analyse particles plugin in Fiji[13]. A ratio of the total area covered by the puncta in the Slit1 producing region to the total area of the forebrain (diencephalon, telencephalon and hypothalamus) was calculated and this value used for statistical analysis.

For the quantification of Sema3A expression in whole brains, a maximum intensity projection was obtained for 60 µm from the z-stack where Slit1 first appears (as Sema3A expression is ventral to Slit1). This maximum intensity projection was used for further analysis. Circular regions of interest (ROIs) were selected in proportion to the size of the brain. One ROI was selected, and one or more background regions were selected in each brain. The ROIs and background regions were selected using the DAPI channel, with ROI being the Sema3A producing region (telencephalon) and background being non-Sema3A producing regions (i.e. the diencephalon and hypothalamus) respectively. The images were manually thresholded (ensuring all RNA puncta were efficiently segmented and the threshold limits were set at the same value for all the images within a replicate) and a binary image was created to segment the RNA puncta. The area covered by the signal was found using the analyse particles plugin in Fiji[13]. Where more than one background region was used, the average area covered by the punctae were used.

$$\text{Normalised expression} = \frac{Total\ area\ covered\ by\ punctae\ in\ region\ of\ interest}{Average\ area\ covered\ by\ punctae\ in\ background}$$

This normalised expression value was used for further statistical analysis. Data were assessed with an unpaired t test with Welch's correction for unequal standard deviation using GraphPad PRISM.

To quantify Sema3A expression in 100 µM LPA-treated embryos, the images were computationally cleared using the Thunder settings in the Leica LAS X software (mounting medium: glycerol, refractive index 1.47290). Thereafter, a maximum intensity projection was obtained for the first 30 µm. The DAPI channel was used to outline the whole brain including the diencephalon, telencephalon, hypothalamus and tectum. Thereafter, the images were thresholded and analysed similarly as described above. The area covered by sema3A punctae was calculated for the whole brain. Normalised expression was calculated by taking a ratio of the area covered by sema3A punctae to the total brain area selected. This normalised expression value was used for statistical analysis. Data were compared using an unpaired t-test with Welch's correction for unequal standard deviation using GraphPad PRISM. PRISM.

To quantify the mRNA expression of *ex vivo* brain regions embedded in hydrogels, the images obtained were randomized and a number was assigned to each image, for blinded analysis. Using the DAPI channel, the tissue regions were first segmented separately for the first 40 µm of the image stacks (i.e. 20 z-slices) using a custom written FIJI script. Additionally, a circular, background, region of interest (5% of image dimensions) was selected where no tissue or collagen hydrogel was present. Mean intensity measurements were then performed in the Sema3A and Slit1 channels for both areas. Subsequently, a median value of the background subtracted signal of interest was calculated in each dissected tissue from all the optical slices. To compare between soft and stiff conditions, mean values were derived independently for each group. Data were assessed using GraphPad PRISM.

For HCR-FISH analysis of the compression stiffening experiments, an image of the embryo was taken after compression stiffening using a CCD camera (Imaging Source, UK) mounted on a TopViewOpticsTM upright imaging system (JPK Instruments AG, Germany). An outline was drawn around the brain, specifically the optic tectum and the bead glued to the cantilever. This was overlayed in Affinity software with the brightfield image of the brain obtained from confocal imaging of the sample on Sp8 Leica Microsystems, laser scanning confocal microscope, to identify the precise region of compression. The region of compression was taken as the ROI (region of interest) and one to two background (BG; non-compressed) regions in the hypothalamus and the diencephalon (caudal to Slit1 expression at the boundary of the telencephalon and diencephalon) were selected. A maximum intensity projection of the first 40 µm was created for both Sema3A and Slit1 and thereafter the images were manually thresholded, segmented, and normalised expression levels were calculated with the same method from the *Sema3A expression in whole brains* section.

***Western Blotting***

Brains of stage 39/40 *Xenopus* embryos were dissected from scrambled morpholino (controls) and Piezo1 morpholino injected embryos. Samples were homogenized in a protease inhibitor cocktail diluted in lysis buffer (NaCl 150 mM, Triton X-100 1%, sodium deoxycholate 0.5%, sodium dodecyl sulfate 4 mg/ml, Tris buffer 50 mM; pH 8), with 1× Halt Protease and Phosphatase Inhibitor Cocktail (Thermo Scientific, UK), and protein samples were prepared for western blot as described previously[14]. Western blotting was done according to LiCor recommendations. Samples were run on 4–12% Bis-Tris NuPage gradient gels (Invitrogen, UK) and transferred to nitrocellulose membrane (Biorad). Membranes were blocked for 30 minutes in a blocking solution of 5% skim milk powder diluted in Tris buffered saline (TBS; pH 7.6), and incubated 1 hour at room temperature with a polyclonal rabbit anti-Piezo1 (anti-FAM38A) primary antibody (NBP1-78446; 1:1,000 dilution), polyclonal rabbit anti-Sema3A antibody (ab199475; 1:500 dilution), monoclonal mouse anti-β-actin antibody (ab6276; 1:1,000 dilution), monoclonal mouse anti-NCAM1 (DSHB Cat# 4d, RRID:AB_528389; 1:2,000 dilution), polyclonal sheep anti-N-cadherin (AF6426-SP; 1:500 diltuion), monoclonal rabbit anti-α tubulin (acetyl K40; ab179484; 1:1,000 dilution), monoclonal mouse anti-α tubulin (ab7291; 1:1,000) in a primary antibody solution of 5% skim milk powder diluted in TBST (TBS+ 0.05% Tween 20; pH 7.6). Excess primary antibodies were then washed off with TBST and the nitrocellulose membrane was incubated for 1 hour at room temperature(18–22°C) in polyclonal goat anti-rabbit antibody conjugated to horseradish peroxidase (HRP) (ab97080; 1:10,000 dilution) for Piezo1,Sema3A and anti-α tubulin (acetyl K40), a polyclonal donkey anti-sheep antibody conjugated to HRP (HAF016; 1:1000 dilution) for N-cadherin, and a polyclonal goat anti-mouse antibody conjugated to HRP (ab6789; 1:10,000 dilution) secondary for β-actin, NCAM1, α-tubulin, in a secondary antibody solution of 5% skim milk powder diluted in TBST (pH 7.6). Western blots were developed using Super Signal West Femto Maximum Sensitivity Substrate ECL (Thermo Fisher Scientific) and the Li-Cor Odyssey FC Imaging system. Signal intensities were measured using the Li-Cor Image Studio Lite software. The ratio of relative intensities of different proteins to total protein staining (Revert 700 total protein stain and wash solution) was used to compare different groups. The average intensities of each protein normalised to total protein stain was calculated for each experiment. To quantify the relative proportion of acetylated α-tubulin in the lysates, the ratio of the relative expression of acetylated α-tubulin to total α-tubulin was obtained.

***Cell density visualization and analysis***

Whole-mount stage 40 *Xenopus* brains with DiI-labelled optic tracts were incubated in 1 µg/mL DAPI (4',6-diamidino-2-phenylindole, Santa Cruz Biotechnology) diluted in 1 × PBS for 10 minutes. Brains were washed thrice in 1 × PBS for 10 minutes and laterally mounted in 1 × PBS for imaging. Z-stacks were acquired using an SP8 confocal microscope (Leica Microsystems, UK; 20× air, NA = 0.75; z-step size = 1 µm). Image stacks were imported into Fiji. For each brain, the image at which the mid-OT bend was in focus was selected. A maximum intensity projection of this image along with the optical section above and below were made. Two 50 µm × 50 µm regions of interest were selected, rostral and caudal to the

OT bend, and a Gaussian blur filter (sigma = 2.0) was applied. Images were manually thresholded to capture all nuclei as accurately as possible. Images were binarised and the 'Analyse Particle' function (size: 1–∞; circularity: 0.2–1.00) was used to acquire the area occupied by nuclei in each region. Relative nuclear density was taken as the ratio of the area occupied by nuclei to the total area of the ROI.

#### *Atomic force microscopy*

##### *Stiffness mapping*

Tipless silicon cantilevers (Arrow-TL1, NanoWorld) were calibrated using the thermal noise method[14] to determine the spring constant *k,* and those with *k* between 0.02–0.04 N/m were selected. Monodisperse spherical polystyrene beads (diameter 37.28 ± 0.34 µm; microParticles GmbH) were glued to the cantilever ends as probes with M-Bond 610 (Agar Scientific). Cantilevers were mounted on a CellHesion-200 AFM head (JPK Instruments), which was set up on an *x*/*y*-motorised stage (JPK Instruments) controlled by custom-written Python scripts[2,15]. Stage 40 embryos (Controls/Piezo1 KD/Sema3A KD/NCAM1 and N-cadherin KD) or stage 35/36 embryos (LPA treated) were anaesthetized with one hemisphere of the brain exposed (see Exposed brain preparation) and transferred to a petri dish on the motorized stage. Indentation measurements were performed automatically in a user-defined grid with a maximum indentation force of 10 nN, approach speed of 5 µm/s (20 µm/s for LPA treated and NCAM1/N-cadherin KD embryos), at a data rate of 1,000 Hz (2,000 Hz for NCAM1/N-cadherin KD embryos). After each measurement, the cantilever was retracted by 100 µm, and the stage moved by a set distance (20–25 µm) to the next position[2,15]. For skin stiffness measurements (Controls/Piezo1 KD), stage 28-31 embryos were anaesthetized (0.04% w/v tricaine methanesulfonate (MS222), 1.3 × MMR, pH 7.6) and immobilized on Sylgard coated dishes (40 mm, TPP). Brightfield and GFP fluorescence overview images were obtained at 25x and 56x magnification with a stereo-fluorescence microscope (AxioZoom.V16, Zeiss) equipped with a Zyla sCMOS camera (Andor) mounted over the AFM setup. Using a custom-written MATLAB script, a grid of AFM measurement points (40 µm × 40 µm) was defined on the skin overlaying the eye primordium, based on the GFP fluorescence of the injected control (scrambled MO)/Piezo1 MO, typically resulting in 21 measurements. Indentation measurements (force setpoint: 2 nN, approach speed: 20 µm/s, data rate: 2,000 Hz).

***Quantification of the reduced apparent elastic modulus K***

Force-distance curves obtained from stiffness measurements were analysed with a custom-written MATLAB script[2,15,19] to obtain the reduced apparent elastic modulus *K*. Raw AFM data were fitted to the Hertz model:

$$F = \frac{4}{3} K \, R^{1/2} \delta^{3/2}$$

where *F* is the applied force, *K* is the reduced apparent elastic modulus $E/(1-v^2)$; where *E* is the Young's modulus and *v* is the Poisson's ratio, *R* the radius of the indenter, and $\delta$ is the indentation depth[20,21].

For exposed brain measurement, force-distance curves were analysed at the maximum applied force *F* = 10 nN. Points where the AFM data were not analysable were excluded. Criteria for excluding individual force-distance curves were (1) inability to apply linear fits through the baseline region, e.g., due to noise, and (2) inability to apply good-quality Hertzian fits to the indentation region, i.e., when the fit aligned poorly with the raw data. The grid of *K* values were plotted as heatmaps in MATLAB, non-analysable measurements were mapped in dark grey. When brain stiffness was calculated by region, a MATLAB script[2] was used to manually select regions of interest using anatomical boundaries and fluorescence-labelling of structures as an approximate guide. For exposed brain measurements, region of interest areas were selected 4 times and pixels defined as part of the region of interest if selected at least 3 out of the 4 times. For skin measurements, force-distance curves were analysed at the maximum applied force *F.* Further to the criteria described above, force-distance curves were excluded from analysis if the maximum applied force deviated by more than 10% from the force setpoint of 2nN. Scripts used for AFM measurements on skin are available from https://github.com/FranzeLab/AFM-data-analysis-and-processing/tree/master/JuliaBeckerThesis .

To determine if the elastic modulus changes during six hour compression stiffening (CS), we compared the median modulus from three force-distance curves acquired in the first hour with the median modulus from three force distance curves acquired in the last hour. The ratio of the median elastic moduli obtained during both time points was calculated for each embryo.

***Compression stiffening***

To induce sustained local compression stiffening[2], tipless silicon cantilevers with $k >$ 0.1 N/m were selected (AFM Probe ARROW-TL1) and polystyrene beads of 89.3-123 µm diameter (microParticles GmbH, Germany) were attached and used to apply a constant force of 30-40 nN in anaesthetized and exposed stage 35/36 embryos (see Exposed brain preparation) where (1) the hypothalamus of wild type embryos or (2) the telencephalon of Piezo1 downregulated embryos, were compressed. The force was applied for > 6 hours at

20-25 °C. Uncompressed controls were treated in the same way except for the AFM application. After every 5-20 minutes of compression, the cantilever was lifted briefly (<10 seconds) to allow for the software to adjust for any drift on the AFM photodiode and a force-distance curve was saved. After removal of the cantilever after > 6 hours, compression-stiffened and control embryos were fixed in modified Carnoy's fixative within 5 minutes and processed for HCR-FISH.

To verify stiffening of brain tissue under uniaxial compression[16] and also to investigate potential changes in tissue viscosity, AFM-based creep measurements were conducted twice on each embryo (Supplementary Fig. 8a). Measurements were carried out with spherical probes of 44.65 µm radius (microParticles GmbH, Berlin, Germany) glued to SICON-TL cantilever (Applied NanoStructures, Inc., Mountain View, CA, USA). Spring constants were determined before the addition of the bead using the thermal noise method implemented in the JPK software and were found to be ~0.18 N/m.

First, indentation measurements were conducted with an extend speed of 10 µm/s and a force setpoint of 10 nN. After reaching the setpoint force $F$ = 10 nN, this force was maintained for 3 seconds, and the tissue's creep response was recorded. Finally, the cantilever was retracted (Supplementary Fig. 8a(left)-b, control experiment *i*). Subsequently, we conducted compression stiffening experiments by applying a force $F$ = 30 nN, which was maintained for 900 seconds (Supplementary Fig. 8a(right), c, CS experiment part *ii*-A). At the end of this force clamp (i.e., of the compression stiffening experiment), the cantilever was moved downward by an additional 10 nN, and another force-hold of 3 seconds at $F$ = 30 nN + 10 nN = 40 nN was performed (Supplementary Fig. 8a (right), c, CS experiment part *ii*-B), in order to assess tissue stiffness at the end of the compression stiffening experiment.

Because of the lack of a baseline in experiment *ii*-B, data could not be fitted with the Hertz model to extract an apparent elastic modulus, *K* (in Pa = N/m$^2$), as in standard indentation measurements. Instead, we fitted a linear function $F = kd + c$ to the initial slopes of the force (*F*) – distance (*d*) curves in approaches *i*, *ii*-A, and *ii*-B to extract a stiffness *k* (in nN/µm) (Supplementary Fig. 8d, e). The stiffness *k* scales linearly with the apparent elastic modulus *K* over a wide range of measurement parameters[15]. In all three approaches, the last 8 nN of the extend segments (before the start of the force clamp) were fitted, and in approach *ii*-A also the initial part of the slope, between 2 nN and 10 nN force application.

To assess the viscosity of the tissue, we fitted a standard linear model, consisting of a spring (with spring constant $k_l$) in series with and element containing a dashpot (with viscosity $\eta$) and second spring (with spring constant $k_a$) in parallel (Supplementary fig. 8f, inset) to indentation-time data. Here, the indentation $\delta$ at time *t* is given as

$$\delta(t) = \left[\frac{3}{4}\frac{F_C}{\sqrt{r}}\alpha^{1-\alpha}\Delta t_a^{\alpha-1}\left(C_0 + (C_0 - C_1)e^{-tC_2}{}_2F_1(\alpha, \alpha+1, \Delta t_a C_2)\right)\right]^{2/3}$$

, where

$$C_0 = \frac{1}{k_l + k_a}, \qquad C_1 = \frac{1}{k_l}, \qquad C_2 = \frac{k_l k_a}{\eta(k_l + k_a)}$$

and r is the radius of the indenter, $\Delta t_A$ the time taken to ramp the force up to the clamp force $F_c$, $\alpha$ is a free parameter describing the shape of the force ramp (1<$\alpha$<2), and ${}_2F_1$ the hypergeometric function. This equation was fitted to the first three seconds of the creep data, i.e., the indentation $\delta$ vs. time *t*, during the force-hold using a custom-written MATLAB script (https://github.com/FranzeLab/AFM-data-analysis-and-processing/tree/master/PilMuk2025).

#### *In vitro techniques*

##### *Single cell dissociation for AFM stiffness measurements*

*Xenopus* embryos at stage 39-40 embryos were anesthetized in MS222 solution (0.04% w/v tricanemethanesulfonate (MS222), 1 × PSF (Penicillin/ Streptomycin/ Amphotericin; Abcam) in 1 × MMR, pH 7.6-7.8) and thereafter immobilised with bent 0.2 mm minutien pins (Austerlitz). The forebrain was dissected using 0.1 and 0.15 mm minutien pins and transferred to a 1.5 mL microcentrifuge tube containing 100 µL of *Xenopus* culture media (1:100 (v/v) dilution of PSF in 6 parts L-15 with 4 parts dd$H_2O$) on ice. Five brains were collected per tube and upon settling at the bottom of the tube, the media was removed. Samples were resuspended in 100 µL of 1 × $Ca^{2+}$- and $Mg^{2+}$-free MMR, with 1% DNase (Roche). The tissues were mechanically dissociated by pipetting them 100 times and incubated at room temperature for 5 minutes before adding 400 µL of *Xenopus* culture media and centrifuging the solution at 400 RCF for three minutes. The supernatant was removed and the pellet was resuspended in 1 mL of *Xenopus* culture media. To functionalize 35 mm glass bottomed dishes (Ibidi), 10 µg/mL poly-D-lysine (PDL, MW 70-150 kDa) in 1 × PBS was added for thirty minutes, followed by 5 µg/mL laminin in 1 × PBS for thirty minutes. Functionalized dishes were washed with *Xenopus* tissue culture media before adding 500 µl of dissociated cell solution and leaving them to adhere at room temperature for 15 minutes, before cellular stiffness was measured with an atomic force microscope.

##### *Single cell stiffness measurements*

Force-distance spectroscopy was performed on a JPK-Bruker CellHesion200 mounted on a Leica DMi8 inverted microscope, with Nanoworld arrow TL-1 cantilevers. The spring constants of the cantilevers were determined using the thermal noise method in the JPK software. Cantilevers with spring constants from 0.044 - 0.1 N/m were selected for this study and a 5.12 µm radius bead was glued to each. The setpoint used to measure cells was 500 pN. Only cells which did not contact other cells and were rounded in appearance (suggesting they had not spread on the glass surface), were measured. Data were processed as described in AFM quantification section. An outlier analysis (ROUT; Q = 1%) was performed using GraphPad PRISM software, and outliers were excluded from further analysis.

***Fabrication of 3D collagen hydrogels***

Preparation of collagen hydrogels was adapted from[16]. Briefly, 25 mM HEPES buffer was dissolved in Leibovitz L-15 medium without L-glutamine and 100 × Penicillin / Streptomycin / Amphotericin (PSF) solution (Abcam) added to achieve a final concentration of 1:100 (v/v). The solution was sterilized with a Nalgene Rapid-Flow PES 0.45 µm filter (Thermo Fisher Scientific), and the mixture was stored at 4°C (for no longer than two weeks). Hydrogel pre-mixes were prepared on ice from 6-parts L15/HEPES/PSF and 1-part (soft gel) or 3 parts (stiff gel) 10 mg/mL bovine collagen stock solution (CellSystems). The remaining volume was filled with autoclaved double-distilled water. Polymerization of collagen hydrogels was initiated by neutralizing the pH to 7.3 with NaOH and leaving the gels at room temperature. After 2 hours, solid gels were overlaid with *Xenopus* culture medium (1:100 (v/v) dilution of PSF in 6 parts L-15 with 4 parts dd$H_2O$) and incubated at 20°C.

***Ex vivo tissue culture in 3D collagen hydrogels***

While pre-mixes of soft and stiff collagen hydrogels were prepared on ice as described above, the hypothalamus and telencephalon brain regions were dissected from stage 37/38 embryos and placed in separate dishes containing *Xenopus* culture medium (1:100 (v/v) dilution of PSF in 6 parts L-15 with 4 parts ddH2O). Around 1 mL of liquid hydrogel solution was then pipetted into a 35 mm glass-bottom dish (Ibidi) and left for pre-polymerization at room temperature for approximately 20 seconds. Around ten brain regions were transferred evenly throughout the solidifying gel. The dish was then kept at 20°C for 2 hours, after which an additional 2 mL of *Xenopus* culture medium was added and brain regions in hydrogels were cultured until 24 hours.

***3D traction force microscopy***

Brain tissue explants from stage 37/38 *Xenopus* embryos were dissected from the hypothalamus region, embedded in soft and stiff collagen hydrogels, and cultured for 24 hours at 20°C (as described in the *ex vivo* tissue culture in 3D collagen hydrogels section). After 24 hours, fresh *Xenopus* culture medium was added. The samples were imaged using an upright confocal laser scanning microscope (TCS SP5, Leica Microsystems) equipped with a 20× water immersion objective (HCX APO L, NA = 1.00, Leica Microsystems). Three image stacks (493 µm × 493 µm × 75 µm) with a voxel size of 481 nm × 481 nm × 2.5 µm, were acquired at 5-minute intervals in reflection mode (pinhole diameter: 2 Airy units), visualising the collagen fibre network, using the 488nm laser line. Following the acquisition of these images, 1 mM Cytochalasin D in DMSO was added to the medium at 1:125 (v/v), and another time-series was acquired with four image stacks (5 minutes apart).

***3D Traction Force Microscopy Analysis***

Analysis of collagen fibre deformations in three-dimensional confocal *z*-stacks was performed using the open-source Python package Saenopy[16]. Stacks acquired prior to Cytochalasin D-induced tissue relaxation were drift-corrected relative to the last stack recorded after Cytochalasin D-treatment. Particle image velocimetry was used for detection of deformation fields using a window size of 30 µm and an element size of 15 µm[16]. Displacement vectors with magnitudes 40% larger than their nearest neighbours were excluded and the resulting fields interpolated to an element size of 10 µm. To solve the inverse problem, material parameters for soft ($k_0$ = 2165, $d_0$ = 0.0022, $\lambda_s$ = 0.0093 and $d_s$ = 0.018) and stiff ($k_0$ = 10076, $d_0$ = 0.0022, $\lambda_s$ = 0.0056 and $d_s$ = 0.027) collagen matrices were used as determined before[16]. For the regularization, an alpha parameter of $10^{8.5}$ with a step size of 0.33 and a fixed number of 150 iterations was used. Displacement and force magnitudes were determined using the Euclidean lengths of individual vectors. Outliers in the reconstructed displacement field were excluded based on a *z*-score threshold exceeding 4, where the *z*-score for each data point was calculated as

$$z = \frac{x - \mu}{\sigma}.$$

Here, *x* represents a single vector magnitude, µ is the mean and *σ* the standard deviation. The strain energy[17] was calculated for the whole stack volume, which corresponds to the total work performed by the tissue to deform its surroundings from the reference state. To reduce variability, scalar quantities were calculated for all vector fields derived from the three image frames of a single explant relative to the last reference frame and the median value determined.

***Statistics***

Names of the statistical tests used, numbers of technical and biological replicates, and *p*-values are provided in the figure captions. All statistical analyses testing a null-hypothesis were two-tailed.

#### Supplementary Figures

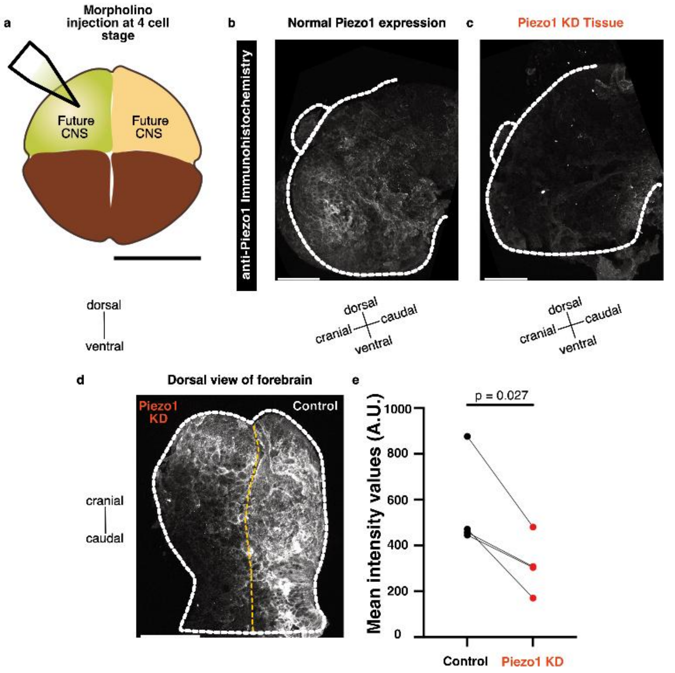


**Supplementary Fig. 1: Downregulation of Piezo1 in half of the developing neuroepithelium. a**, Schematic. At the 4-cell stage, translation-blocking morpholinos were injected into one of the two dorsal (usually less pigmented) blastomeres, which contribute to the formation of the nervous system. Scale bar: 600 µm. **b-d,** Immunofluorescence images of *Xenopus laevis* brains at stage 40. The dashed white outlines depict the boundaries of the brain tissue. Scale bars: 100 µm. **b-c,** Piezo1 expression patterns in neuroepithelium (lateral view) arising from the **b,** uninjected blastomere and **c,** the blastomere injected with translation-blocking Piezo1 morpholino, resulting in Piezo1 knockdown (KD). **D,** Dorsal view of Piezo1 expression in the developing neuroepithelium showing both the injected (Piezo1 KD) and the uninjected (Control) hemispheres. The dashed yellow line represents the midline. **E,** Piezo1 morpholino injection into one dorsal blastomere led to a significant decrease in Piezo1 expression in one half of the developing neuroepithelium. The graph represents the paired (Control vs Piezo1 KD) mean intensity values of the developing telencephalon. Each point represents an embryo (*N* = 4). Data was assessed with a ratio paired t-test; *p*-value indicated in the figure. KD: knockdown.

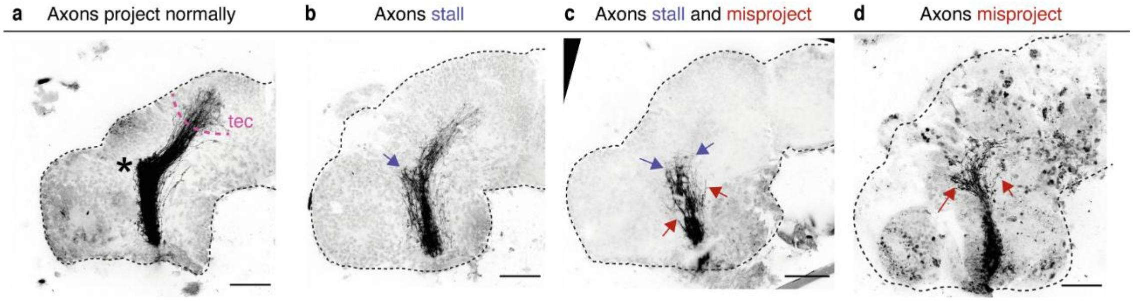


**Supplementary Fig. 2: RGC axon growth phenotypes and guidance defects in the developing *Xenopus* brain. a**, Normal RGC axonal projections in the brain of a stage 40 *Xenopus laevis* embryo. RGC axons form a bundle, turn in the mid-diencephalon (marked with '*'), and arrive at the optic tectum (tec). **b,** Representative image of a brain showing stalling of some RGC axons (purple arrow) before others turn caudally in the mid-diencephalon. **c,** Representative image of a brain showing RGC axon stalling (purple arrows) and splaying (i.e., unbundling / defasciculation), leading to misprojections (red arrows). **d,** Representative image of a brain showing RGC axons unbundling and misprojecting (red arrows). Scale bars: 100µm. RGC: retinal ganglion cell.

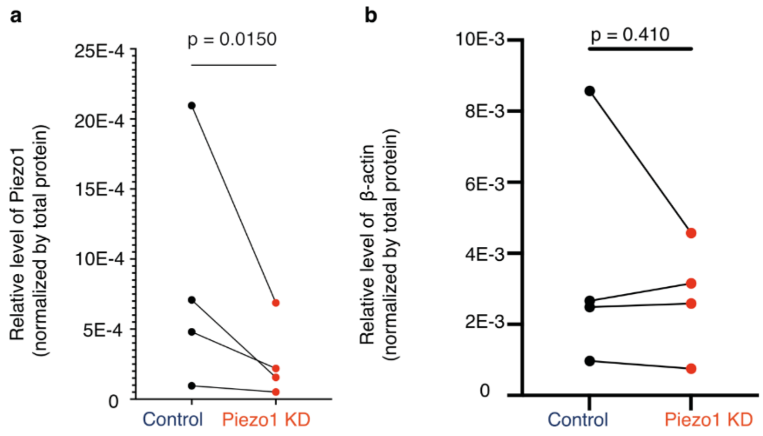


**Supplementary Fig. 3: Quantification of Western blots.** Downregulating Piezo1 led to a significant decrease in the expression of Piezo1 protein **a,** but not of β-actin **b**. Representative Western blot images are shown in Fig. 2k. Data were normalised by the total protein concentration. Each point represents the mean of a biological replicate. Data were assessed with a ratio paired t-test. *p*-values are indicated in the figure. KD: knockdown.

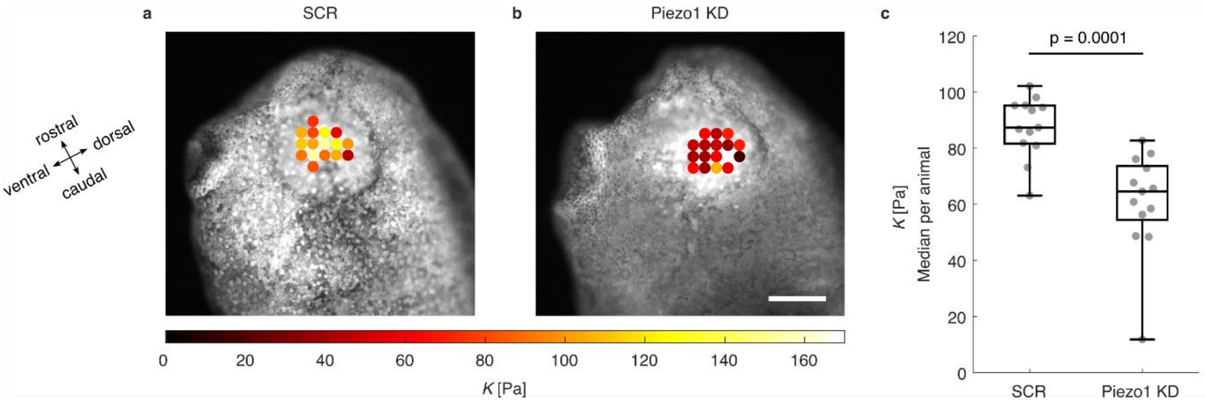


**Supplementary Fig. 4: Piezo1 knockdown leads to softening of skin tissue. a, b,** Exemplary heatmaps of AFM measurements for stage 31 **a,** SCR (control) and **b,** Piezo1 knockdown (KD) embryo skin tissue, overlaid on a fluorescence image of the GFP-containing injected constructs. Heatmaps are scaled to an apparent elastic modulus of 170 Pa; scale bar is 200 µm. **c,** Median skin tissue stiffness per animal in both conditions. Each dot corresponds to one animal; boxes denote 1st to 3rd quartiles with annotated median, whiskers extend from min to max. Both groups were compared with a nested t-test, where AFM measurements were nested under animals, and animals under condition. N = 13 animals per condition; n = 3 - 23 (mean: 16.5) stiffness measurements per animal.

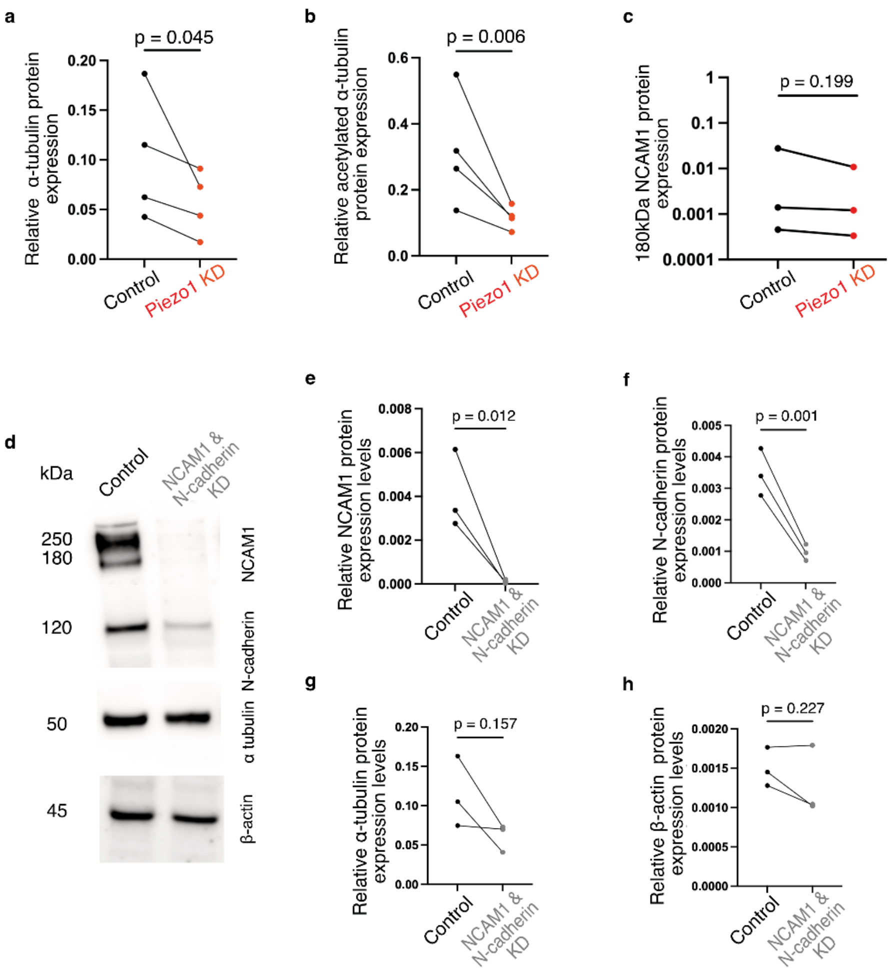


**Supplementary Fig. 5: Cytoskeletal changes following genetic perturbations. a-c**, Downregulating Piezo1 led to decreased expression of α-tubulin **a,** and acetylated α-tubulin **b**, but not of 180 kDa NCAM1. **c**. Representative Western blot images are shown in Fig. 4a. Data were normalised by the total protein concentration. Each point represents the mean of a biological replicate. Data were assessed with a ratio paired t-test. **d,** Representative Western blot of NCAM1, N-cadherin, α-tubulin and β-actin expression in control and NCAM1 and N-cadherin-knockdown samples (KD). **e-h**, Morpholino-mediated knockdown of NCAM1 and N-cadherin led to decreased protein expression of **e**, NCAM1, and **f**, N-cadherin but not of **g**, α-tubulin and **h**, β-actin. Data were normalised to total protein levels and compared with a ratio paired t-test. KD: knockdown.

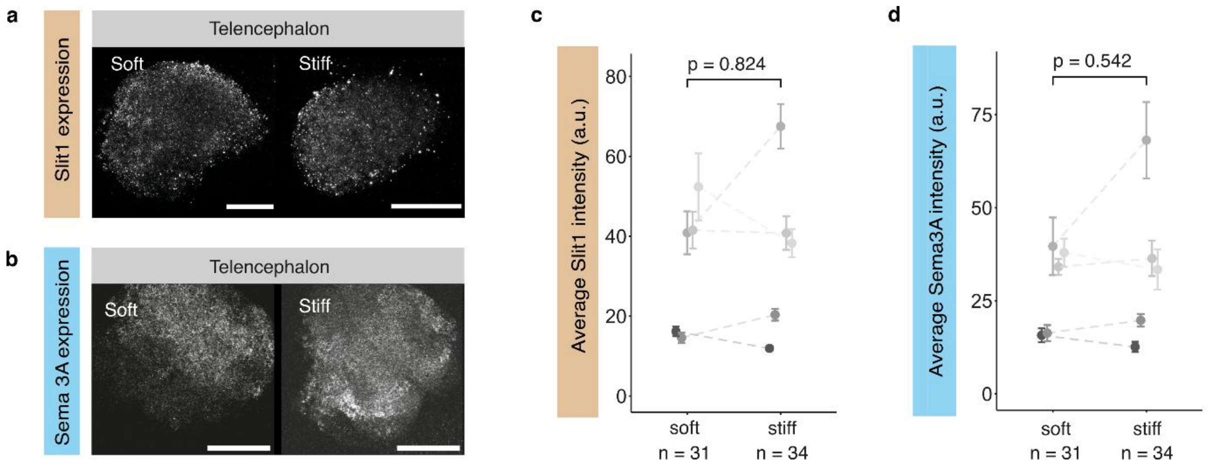


**Supplementary Fig. 6: Chemical cue expression in the telencephalon is independent of substrate stiffness *ex vivo*.** **a-b,** Representative HCR-FISH images of **a**, Slit1 and **b**, Sema3A expression in telencephalic tissue after 24 hours in soft (left) and stiff (right) substrates. Scale bars: 100 µm. **c-d,** Quantification of mRNA expression. In telencephalic tissue, neither **c**, Slit1 nor **d**, Sema3A expression changed with substrate stiffness. Individual points represent means with standard errors of each biological replicate. A ratio paired t-test was used for statistical testing. *p*-values are indicated in the graphs. a.u.: arbitrary units. n denotes the number of tissue explants from 5 independent experiments.

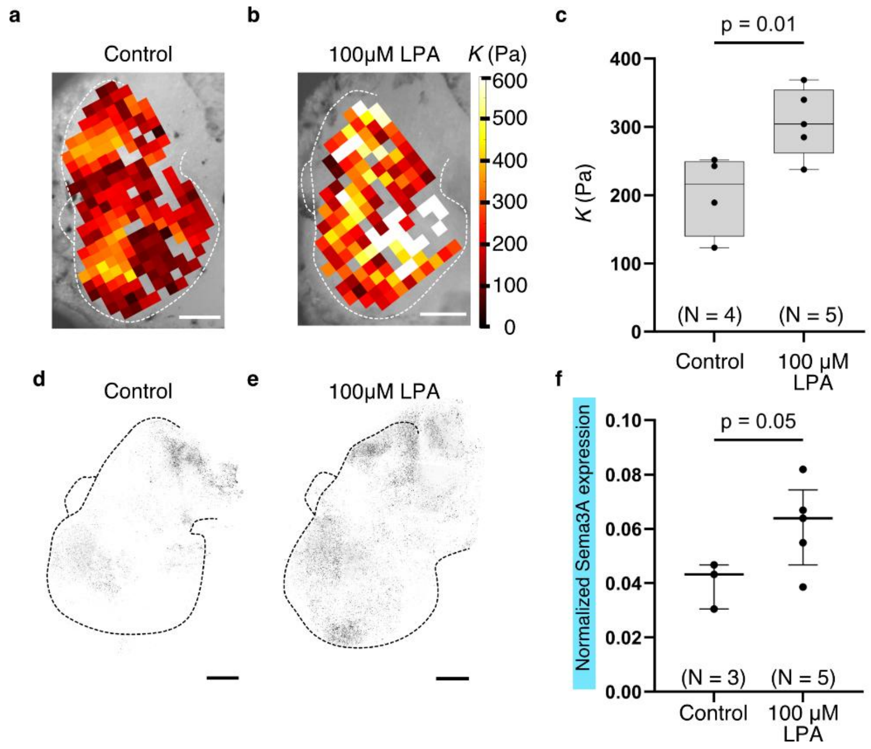


**Supplementary Figure 7: Lysophosphatidic acid (LPA) treatment results in increased tissue stiffness and Sema3A expression. a-b,** Representative heatmaps of Stage 33/34 embryonic brains treated for 2-3 hours with either **a,** 100 µM DMSO (control) or **b,** 100 µM LPA, overlayed on a brightfield image of the brain. The brain boundary is outlined by a white dotted line. Heat maps are scaled to an apparent elastic modulus *K* = 600 Pa. **c,** Box plots of median apparent elastic moduli *K* per animal in both conditions. Each dot in the graph corresponds to an animal measured. The boxes denote the $1^{st}$-$3^{rd}$ quartiles with a line representing the median. The whiskers extend from min to max. Groups were compared with a nested t-test, where AFM measurements were nested under animals, and animals under condition. *K* increased significantly in LPA-treated brains as compared to controls (*p* value indicated in the Figure). **d-e,** Representative images of Sema3A expression in embryonic brains treated for 6 hours with **d,** 100 µM DMSO (control) or **e,** 100 µM LPA. The brain boundary is outlined by a black dotted line. **f,** Scatter dot plot showing normalized Sema3A mRNA expression levels. Each dot corresponds to an embryo measured.

Normalized Sema3A expression (calculated as the ratio of the area occupied by Sema3A puncta in the whole brain to the total brain area) significantly increased upon 100 µM LPA treatment for 6 hours as compared to controls (treated with 100 µM DMSO). Data were compared with a two tailed unpaired t-test with Welch's correction (*p* value is indicated in the figure). Scale bars: 100 µm. N: number of embryos measured.

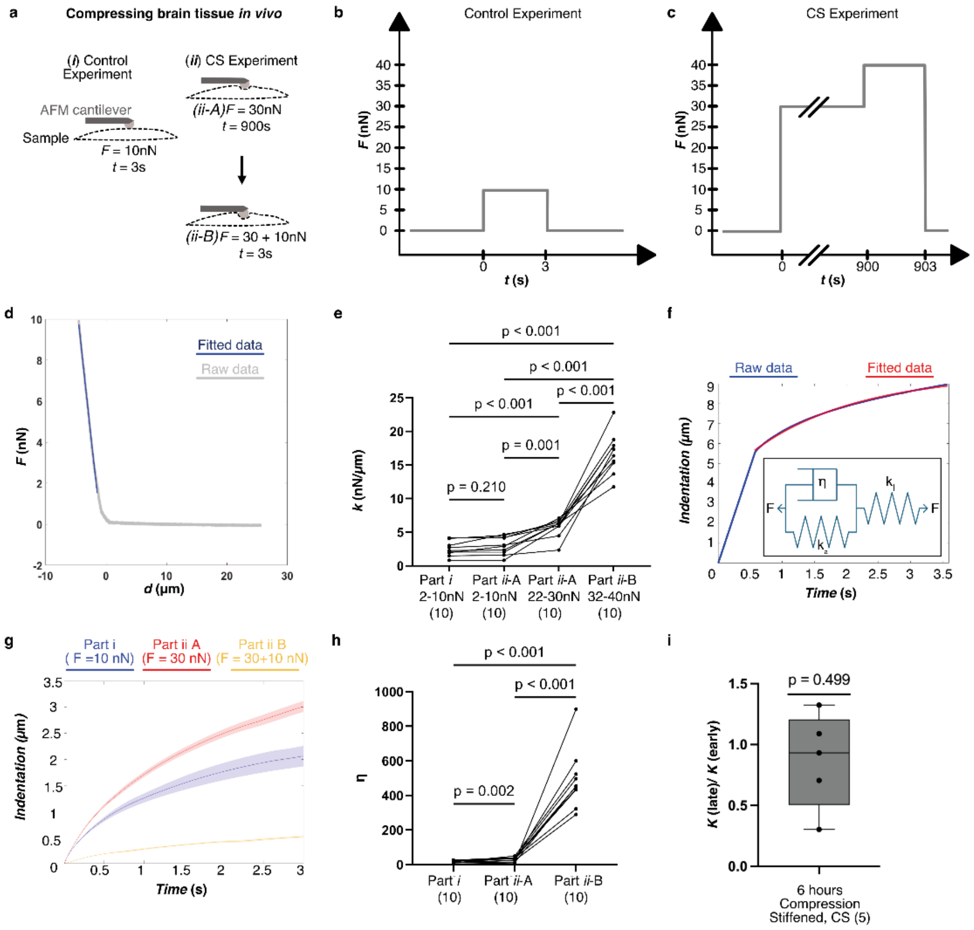


**Supplementary Figure 8: Uniaxial compression leads to brain tissue stiffening *in vivo*.** **a-c**, Schematic of the experiment. In the experiment mimicking the control condition (**a***(i)*, **b**), the tissue was indented with *F* = 10 nN for a duration of *t* = 3 s. In the experiment mimicking the compression stiffening (CS) experiments (**a***(ii)*, **c**) (Fig. 6), the tissue was first exposed to a constant force of *F* = 30nN for 900 seconds (*ii*-A), and subsequently, without retracting the cantilever, it was indented further by an additional *F* = 10nN for 3 seconds (*ii*-B). **d**, Example of a linear fit (blue) to a force-distance (*F-d*) curve (grey) used to calculate the stiffness *k* (see Methods). **e**, Plot of the stiffnesses obtained by fitting $F = kd + c$ through the last 8 nN of the *F-d* curves of indentations (*i*), (*ii*-A), and (*ii*-B), as well as through the range of 2 nN – 10 nN of indentation (*ii*-A). The stiffness of the tissue was similar during the first 10 nN indentation in approaches (*i*) and (*ii-A*), as expected. However, already during the application of the compressive force in approach (*ii-A*), tissue stiffness increased immediately. $k$ was significantly smaller at the beginning (2-10 nN) of the indentation than at its end (22-30 nN indentation) (median $k_{2\text{-}10nN}$ = 3.0 nN/µm *vs.* $k_{22\text{-}30nN}$ = 6.2 nN/µm). When probing tissue stiffness after compression for 900s (*ii-B*), $k$ increased even further (median $k_{ii\text{-}B}$ = 16.9 nN/µm). These data confirmed that uniaxial compression leads to tissue stiffening[18] *in vivo*. As the developing *Xenopus* brain is fairly homogeneous at early stages of development[19], an impact of potentially stiffer, deeper layers to the measured increase in stiffness it rather unlikely. **f,** Example indentation-time ($\delta$-$t$) curve (blue) with a Standard Linear Model (SLM) fit (red); see Methods. Inset: schematic diagram of the SLM, which consists of a spring in series with an element containing a parallel dashpot (with viscosity $\eta$) and a second spring. **g**, Mean indentation-time ($\delta$-$t$) curves for (*i*), (*ii-A*) and (*ii-B*), with the shaded regions representing +/- the Standard Error of the Mean. The creep (flow) of the tissue was relatively smaller for larger forces. **h**, Quantification of the apparent viscosity $\eta$. $\eta$ increased significantly with increasing forces, indicating that the tissue became less fluid under compression, in particular on longer time scales. **e**, **h**, Data were compared with a repeated measures one-way ANOVA with Geisser-Greenhouse correction, followed by a Tukey's multiple comparison test. **i**, The ratio of the median apparent elastic modulus $K$ obtained in the last hour of the > 6 hour compression stiffening experiment to that obtained in the first hour was similar to 1 (one sample $t$-test), suggesting that tissue mechanics was only altered for as long as compression was applied. Each data point represents an embryo measured. Numbers in parentheses indicate the number of measurements. The boxes span the first and third quartiles with a line at the median. The whiskers span the range of the data. All $p$-values are indicated in the figure. CS: compression stiffening.

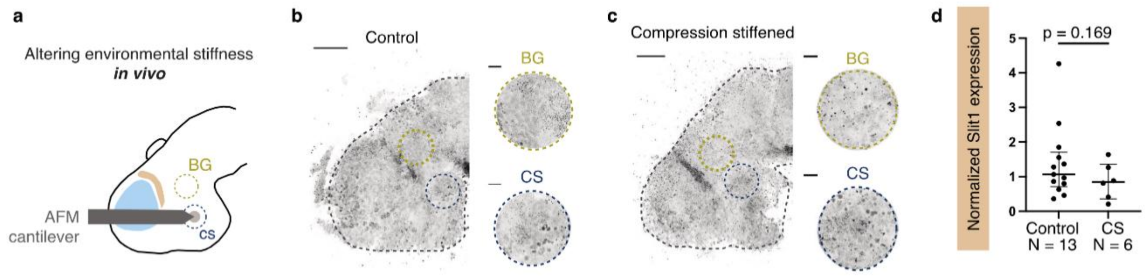


**Supplementary Figure 9: Compression-stiffening of hypothalamic tissue *in vivo* does not significantly alter Slit1 expression. a,** Schematic representation of the experimental setup for compression stiffening brains *in vivo*. **b-c,** Representative images of Slit1 expression in **b**, control and **c**, compression-stiffened brains. Dashed circles indicate compression-stiffened (CS) and background (BG) regions used for analysis. The insets show a magnified view of representative areas. Scale bars: 100 µm for whole brain images, 20 µm for insets. **d,** Normalised Slit1 expression levels (calculated by measuring the ratio of the total area covered by the signal in the CS region to the mean area covered by the signal in the BG regions) did not significantly differ between compression-stiffened brains and controls. Individual dots represent the normalised expression levels in each embryo. Bars represent the median and the interquartile range. An unpaired t-test with Welch's correction was used for statistical analysis. The p value is indicated in the figure. BG: background, CS: compression stiffened.

**Supplementary Videos**

**Supplementary Video 1: Long-term physical interactions between a brain tissue explant and a soft hydrogel matrix.** Brain tissue from the hypothalamic region of a developing *Xenopus* brain was cultured in a soft collagen hydrogel for 24 hours. Maximum intensity-projected time-lapse acquisition (5 minutes between frames) of confocal stacks in reflection mode revealed dynamic matrix deformations at the tissue – fibre network interface. Scale bar: 40 µm.

**Supplementary Video 2: Traction force microscopy of brain explants in hydrogel matrices.** Brain tissue explants were cultured in soft and stiff collagen hydrogels for 24 hours. Three confocal reflection stacks were acquired every 5 minutes prior to inducing tissue relaxation by the application of Cytochalasin D, and another four stacks acquired immediately after. Here, the maximum intensity projection of a hypothalamic tissue explant in a soft collagen hydrogel is shown. White arrows indicate neurites which extended from the explant; the dashed white line marks the tissue boundary. The 3D displacement field was calculated between the first and last image stack and sum-projected along the z-axis to obtain a 2D representation with colour indicating the vector magnitudes. Scale bar: 150 µm.
